# Motor Learning and Transfer Are Symmetric Across Hands

**DOI:** 10.64898/2026.07.17.739258

**Authors:** Indranil Nyamsuren, Ashley Statham, Elise Mitchell, Brayden Kohler, Phoebe Lam, Roberta L. Klatzky, Jonathan S. Tsay

## Abstract

Functional asymmetry between the cerebral hemispheres is a defining feature of the sensorimotor system, with the dominant hemisphere playing a central role in *motor control*. Whether *motor learning* is similarly lateralized, however, remains unresolved. To tackle this question, we combined a comprehensive meta-analysis (114 datasets) with a series of well-powered, preregistered experiments (N = 526) to test two core behavioral predictions of hemispheric lateralization in sensorimotor adaptation, a canonical form of motor learning: (1) adaptation is preferentially expressed in the dominant hand and (2) transfers asymmetrically between limbs. Across both approaches, we found that adaptation and interlimb transfer were strikingly symmetric. Together, these findings support a fundamental dissociation in the neural organization of skilled behavior: whereas motor control is lateralized to the dominant hemisphere, motor learning is supported by a neural architecture that functions symmetrically.

## Introduction

The human brain is known to be asymmetric, with many core functions specialized to one hemisphere^1–16^. Classic neuropsychology provided some of the earliest and most compelling demonstrations of this principle. In the domain of language, Paul Broca’s observations of aphasia following left-hemispheric lesions^17,18^ led him to conclude that “we speak with the left hemisphere”^19^, a claim later substantiated by anatomical and neuroimaging evidence^20–22^. Conversely, spatial attention has long been associated with the right hemisphere, where lesions often give rise to hemispatial neglect^23,24^. Neuroimaging^25–31^, together with comparative evidence across species^32,33^, has reinforced these observations, converging on the view that hemispheric specialization is a defining principle of the human nervous system.

Whereas hemispheric specialization in cognitive domains is often framed as a left–right dichotomy, specialization in the motor system is organized around the dominant hemisphere, which plays a central role in **motor control**^34–40^. This functional asymmetry underlies handedness^41–43^, with skilled movements of the dominant hand controlled primarily by the contralateral dominant hemisphere^41–43^. Consistent with this specialization, the dominant hemisphere exhibits greater corticospinal excitability^41–43^, more efficient sensorimotor processing^44–46^, and stronger functional connectivity within motor networks^47,48^ than its nondominant counterpart.

Whether this same organizing principle extends to **motor learning**, however, remains far less clear^49,50^. A longstanding hypothesis is that motor learning, like motor control, is specialized in the dominant hemisphere^51,52^. This hypothesis has been studied most extensively through sensorimotor adaptation (sensorimotor learning)—the process by which the nervous system learns from sensory prediction error, the mismatch between predicted and actual sensory consequences of movement. According to this view, the dominant hemisphere generates more accurate sensory predictions, enabling greater learning when those predictions are violated^51–53^. Because transforming sensory prediction errors into improved future movements is a core component of motor learning^54^, sensorimotor adaptation has become the principal paradigm for investigating its hemispheric organization.

The dominance account of learning makes two key predictions. First, by the preceding arguments, adaptation should be greater in the dominant than the nondominant hand; that is, motor learning should be asymmetric across limbs. Second, adaptation should transfer asymmetrically between the limbs, with greater transfer toward than away from the dominant hand^55–59^. This latter prediction follows from the assumption that given its stronger coupling across sensorimotor cortices^47^, the dominant hemisphere learns from errors generated by the ipsilateral hand to a greater extent than the nondominant hemisphere, which primarily learns from errors generated by the contralateral hand it more directly controls.

Despite the conceptual simplicity of these hypotheses, the literature has yielded remarkably inconsistent findings in their support. For the first prediction, some studies report a dominant-hand advantage in motor adaptation^60–63^, whereas others find no difference between hands^64,65^ or even superior adaptation in the non-dominant hand^66,67^. Evidence for the second prediction is similarly mixed: some studies report greater transfer toward the dominant hand^56,68–70^, others report greater transfer away from the dominant hand^36–38^, and still others symmetric interlimb transfer^71^. Consequently, a fundamental question remains unresolved: **Does motor learning follow the same principles of hemispheric specialization that govern motor control, such that the dominant hemisphere enjoys an advantage in both learning and control?**

Two methodological limitations may help explain these inconsistencies. First, studies testing the effects expected from hemispheric specialization may have been underpowered to detect the corresponding behavioral differences^72^. Second, classic adaptation paradigms fail to isolate implicit motor adaptation—the error-driven sensorimotor process most directly implicated by the dominant-hemisphere hypothesis—and instead permit substantial contributions from explicit strategy use^66,73^. Consequently, contamination from these cognitive strategies may have obscured subtle asymmetries in implicit sensorimotor adaptation and interlimb transfer, masking the putative signatures of hemispheric specialization for motor learning.

To address these limitations, we adopted a two-pronged approach: First, we conducted a comprehensive meta-analysis to re-examine whether motor adaptation is preferentially expressed in the dominant hand and whether it transfers asymmetrically between the limbs (114 datasets). Second, we performed a series of preregistered, well-powered experiments in both left- and right-handed participants using paradigms of motor adaptation designed to dissociate explicit and implicit learning processes (N = 526). Specifically, implicit motor adaptation was isolated by a task-irrelevant visuomotor perturbation that the nervous system interpreted as a relevant motor error, despite explicit knowledge to the contrary^74,75^.

These approaches enabled us to distinguish among three competing theories of the lateralization of learning in relation to hemispheric organization: Under a **dominance-advantage account** (Fig. 1A), in which the hemisphere controlling the dominant hand is specialized for learning, adaptation should mirror motor performance, such that greater adaptation is found in the dominant hand irrespective of handedness^52,76^. Under a **left-advantage account** (Fig. 1B), whereby learning is lateralized to the left hemisphere, adaptation should be greatest in the right hand irrespective of handedness^8^. Critically, because most studies have exclusively studied right-handers, they cannot distinguish whether apparent learning advantages in the right hand reflect specialization of the left hemisphere or the hemisphere controlling the dominant hand. Left-handers provide the critical test between these alternatives: if learning follows hand dominance, the advantage should reverse to the left hand; if it reflects left-hemisphere specialization, the right hand should retain its advantage. Finally, under a **functional-symmetry account** (Fig. 1C), both hemispheres possess equal capacity for learning, yielding equivalent adaptation and interlimb transfer^77–81^. We return to the neural implications of these competing accounts in the Discussion.

**Figure 1.**
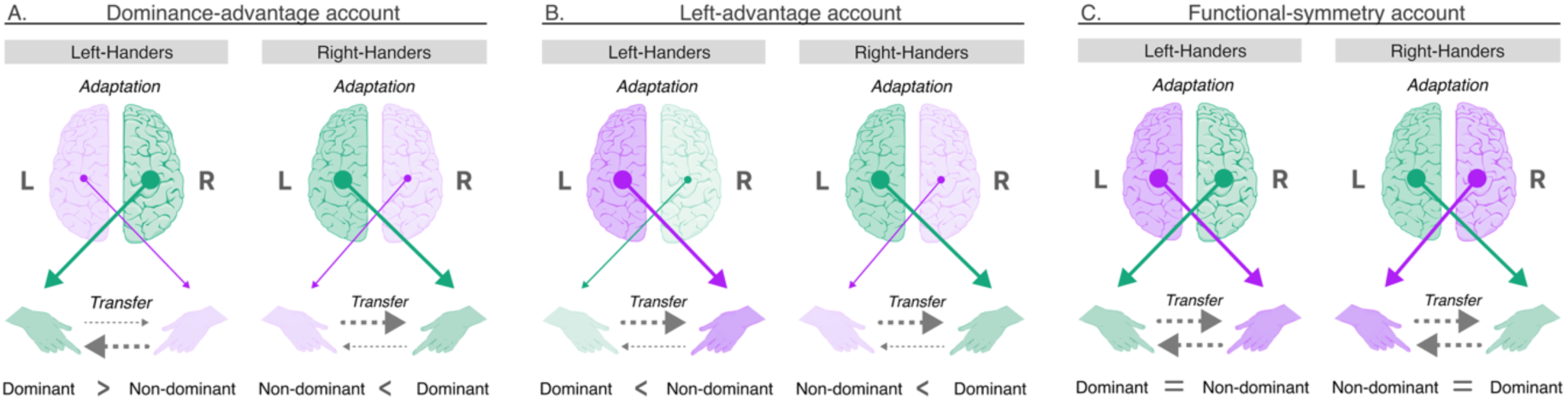
Competing Accounts of Hemispheric Organization of Motor Learning. **(A)** The dominance-advantage account predicts greater implicit motor adaptation in, and greater transfer towards, the dominant hand regardless of handedness. **(B)** The left-advantage account predicts greater implicit motor adaptation in, and greater transfer towards, the right hand regardless of handedness. **(C)** The functional-symmetry account predicts equal implicit motor adaptation and interlimb transfer across both hands. Colored arrows indicate the magnitude of learning contributed by the left and right hemispheres, expressed through their respective contralateral hands. Gray arrows depict the magnitude of transfer toward and away from the dominant hand. Larger arrows indicate greater adaptation and/or interlimb transfer.

Furthermore, whether explicit strategy use exhibits comparable behavioral asymmetries in motor learning remains unknown. This distinction is theoretically important because explicit and implicit learning are thought to rely on different computational mechanisms. Whereas implicit adaptation has been hypothesized to arise from effector-specific sensorimotor processes, explicit strategy use depends on top-down cognitive representations that can presumably be flexibly deployed to either hand. Consequently, explicit strategy use is predicted to exhibit little or no asymmetry, consistent with a functionally symmetric neural organization.

## Results

### Meta-analysis 1: No Evidence for a Dominant-Hand Advantage in Sensorimotor Adaptation

We began by asking whether motor learning exhibits the hallmarks of hemispheric specialization. To provide the most comprehensive assessment to date, we conducted the first meta-analysis of studies examining whether learning is asymmetrically expressed in the dominant hand and whether it transfers asymmetrically between the limbs. Following the literature, we focused our synthesis on motor adaptation because it provides a well-characterized model of motor learning^54^.

We identified 46 datasets derived from 23 studies that met our eligibility criteria (Fig. 2A). Reflecting sustained interest in this question, these studies span research conducted over the last two decades (Fig. 3C) and collectively include 635 participants (Fig. 3E). The majority of studies employed visuomotor perturbation tasks, in which participants adapted their movements to compensate for a rotation of the visual cursor relative to the hand, thereby restoring accurate performance. Other studies employed prism adaptation tasks, in which prism goggles displaced visual feedback during goal-directed movements, or force-field adaptation paradigms, in which external forces systematically perturbed reaching trajectories and participants learned to counteract these dynamic disturbances (Fig. 3A).

**Figure 2.**
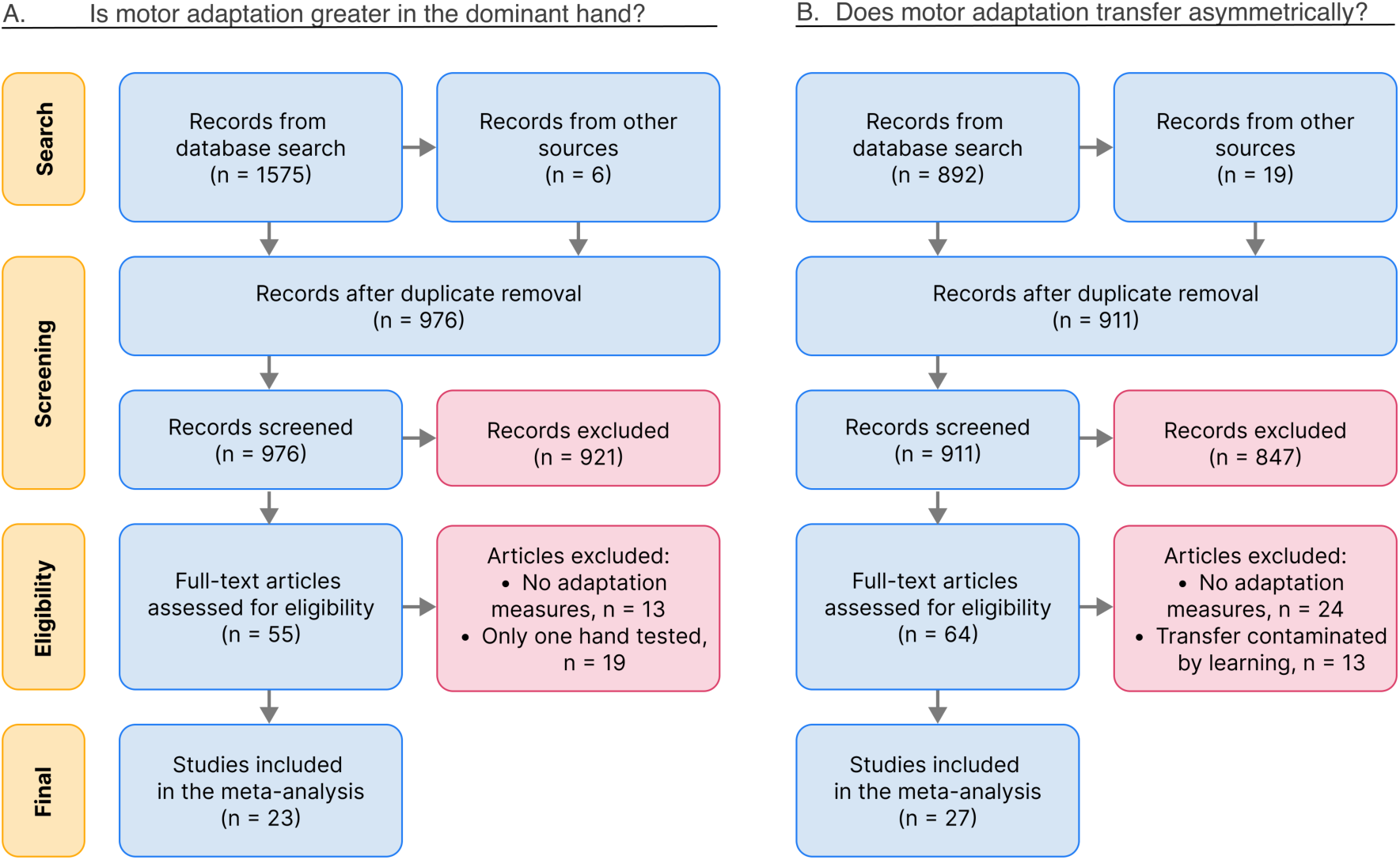
PRISMA Flow Chart. **(A)** The first meta-analysis examined whether sensorimotor adaptation is preferentially expressed in the dominant or non-dominant limb (46 eligible datasets derived from 23 studies). **(B)** The second meta-analysis examined whether sensorimotor adaptation transfers asymmetrically between limbs (68 eligible datasets derived from 27 studies).

**Figure 3.**
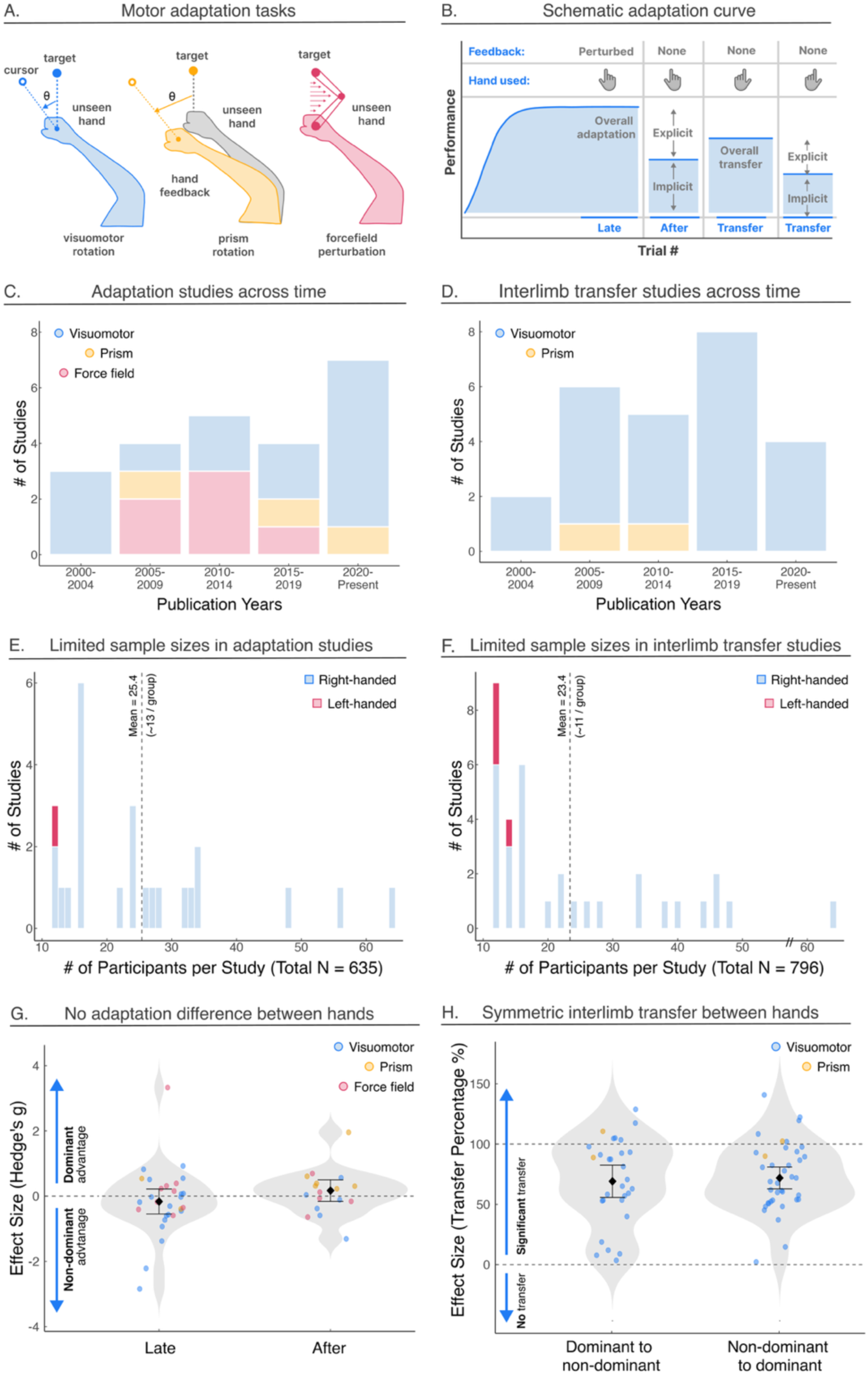
Meta-analysis Overview. **A)** Common sensorimotor adaptation tasks in the literature included visuomotor rotation, prism adaptation, and force-field adaptation. **B)** Schematic of behavior in a standard motor adaptation task. Following a baseline block (not shown), a perturbation is introduced. Performance during the late adaptation phase provides a measure of overall learning, reflecting the combined contributions of implicit adaptation and explicit strategy use. In the subsequent no-feedback aftereffect block, the perturbation is removed, and participants are instructed to forgo any re-aiming strategy and reach directly to the target. Because explicit strategies are minimized, performance during this phase provides a selective measure of implicit adaptation. During the interlimb transfer phase, participants switch to the untrained hand and perform reaches without visual feedback. Transfer can be assessed either by instructing participants to maintain their learned aiming strategy, yielding a measure of overall transfer that includes both implicit and explicit components, or by instructing them to reach directly to the target, yielding a more selective measure of implicit transfer. **C-D)** Number of studies by publication year. **E-F)** Sample sizes included in both meta-analyses. **G)** No significant differences in motor adaptation between dominant and non-dominant hands. Translucent dots represent individual datasets; colors indicate task type. The solid dot denotes the grand effect size; error bars indicate the 95% confidence interval. Negative effect sizes indicate greater adaptation in the dominant limb, whereas positive effect sizes indicate greater adaptation in the non-dominant limb. **H)** Interlimb transfer was significant and symmetric toward vs away the dominant limb. Horizontal dashed lines at 0% and 100% denote no transfer and full transfer, respectively.

We focused on two commonly reported measures of motor adaptation: The first was *late adaptation*, defined as performance after prolonged exposure to a perturbation. Because late adaptation reflects the combined contributions of implicit and explicit learning, it provides a measure of overall adaptation but cannot distinguish between these processes^54^ (Fig. 4B). The second was the *aftereffect*, measured once the perturbation had been removed. Persistent deviations in movement direction during this post-learning phase reflect the lingering effects of adaptation and provide a more selective measure of implicit adaptation^54^. For both measures, evidence of hemispheric specialization would be reflected in a consistent learning advantage in one hand^51,52^.

**Figure 4.**
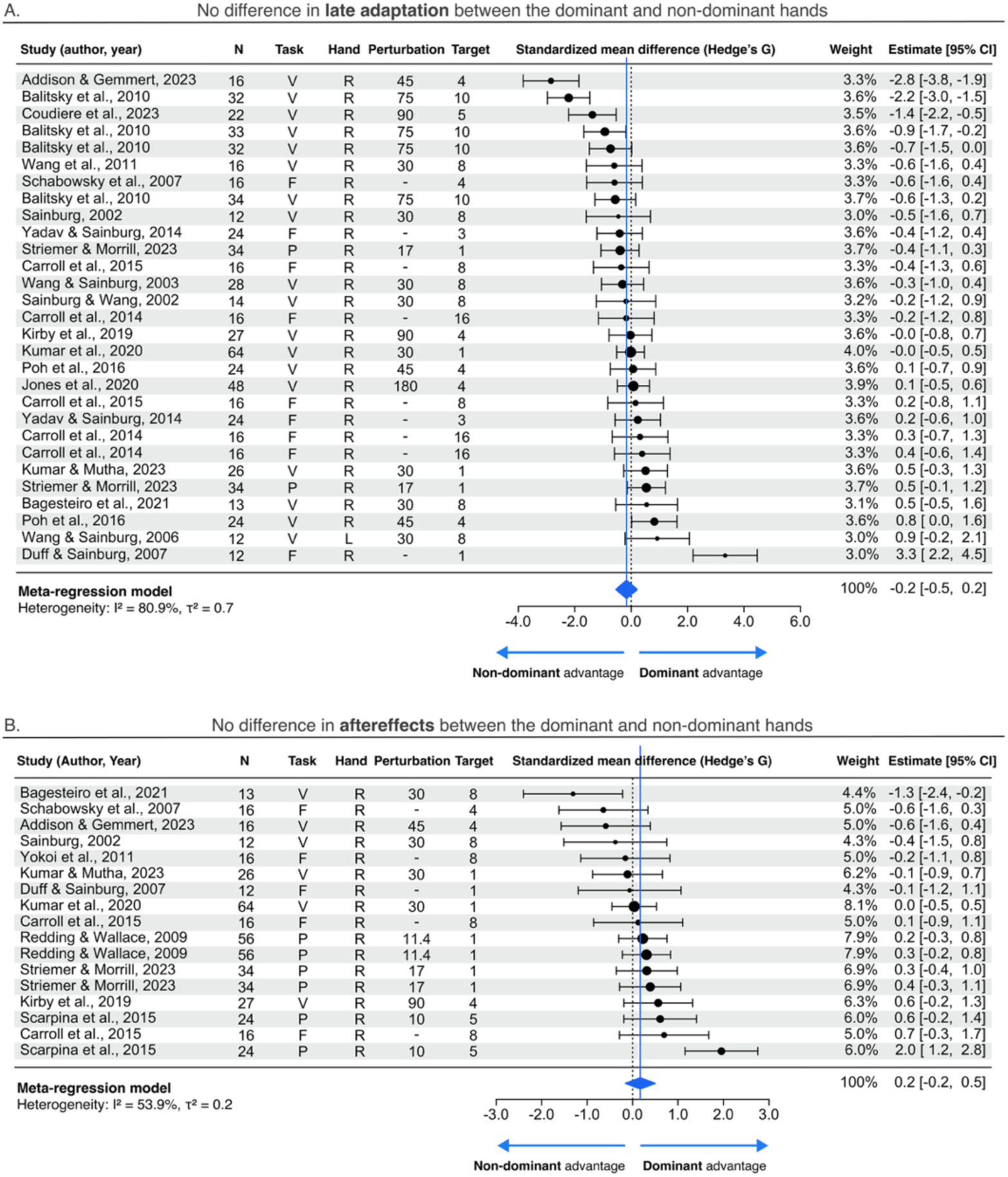
No significant difference in motor adaptation between dominant and non-dominant hands. **A-B)** Forest plots aggregate **A)** 29 datasets quantifying late adaptation and **B)** 19 datasets quantifying aftereffects. The diamond denotes the overall effect size (Hedges’ g) and its 95% confidence interval. The N column shows the number of individuals in the dominant (D) and non-dominant (ND) groups. The size of each effect-size marker reflects the weight assigned to that study in estimating the pooled effect size. In the Task column, V denotes visuomotor adaptation, P prism adaptation, and F force-field adaptation. The Hand column shows the self-reported handedness of participants. The Perturbation column indicates the perturbation magnitude (°) used in visuomotor and prism adaptation tasks. The Target column shows the number of training targets. Negative effect sizes indicate greater adaptation in the dominant limb, whereas positive effect sizes indicate greater adaptation in the non-dominant limb.

Twenty-nine datasets assessed the effects of hand dominance on *late adaptation*. Effect sizes varied widely across studies (*τ*^2^ = 0.7, *I*^2^ = 80.9%) ranging from a Hedge’s *g* of -2.8 (a negative effect denotes greater late adaptation in the non-dominant hand) to 3.3. Despite this heterogeneity, the pooled effect size was not significantly different from zero, providing no evidence for a systematic difference in late adaptation between dominant and non-dominant hands (Fig. 3G & Fig. 4A; b = -0.1 ± 0.2, t(18.8) = −0.4, p = 0.7).

A similar pattern emerged for *aftereffects*. Across 17 datasets, effect sizes ranged from *g* = -1.3 to 1.9 and again exhibited significant heterogeneity (*τ*^2^ = 0.2, *I*^2^ = 53.9%). Nevertheless, the pooled effect size did not differ significantly from zero (Fig. 3G & Fig. 4B; b = 0.04 ± 0.2, t(11.3) = 0.3, p = 0.7, indicating no systematic difference in implicit adaptation between dominant and non-dominant hands. When considered alongside the null effects for late adaptation (*τ*^2^ = 0.5, *I*^2^ = 74.2%; b = -0.06 ± 0.2, t(21.7) = -0.4, p = 0.7), these findings underscore the absence of a dominant-hand advantage in both the implicit and explicit components of motor adaptation.

Importantly, these null effects were robust across experimental contexts. Effect sizes did not differ across different types of motor adaptation paradigms (visuomotor: b = 0.1 ± 0.2, t(5.9) = 0.9, p = 0.4 vs. prism: b = 0.01 ± 0.2, t(7.9) = 0.05, p = 0.9 vs. force-field: b = -0.4 ± 0.2, t(2.9) = 1.7, p = 0.2). Nor did they vary as a function of perturbation size in visuomotor tasks (range of rotations: 10–180°; b = -0.002 ± 0.005, t(2.7) = -0.3, p = 0.7) or the number of movement targets (range: 1–16°; b = -0.0004 ± 0.005, t(2) = -0.09, p = 0.9). We did not find any evidence for publication bias (b = -0.2 ± 1.4, t(9.2) = -0.1, p = 0.9). Thus, across a broad range of experimental results, we found no evidence for either a dominant- or non-dominant-hand advantage in motor adaptation.

The literature was characterized by relatively small sample sizes, with an average of 25 participants per study (approximately 13 participants per group). As a consequence, statistical power was low (median statistical power [IQR]: 5.2% [5.1%, 5.3%], with α = 0.05). Another major limitation was the near-exclusive focus on right-handed participants. With the exception of a single dataset^55^, all studies were conducted exclusively with right-handers, making it difficult to distinguish whether learning advantages reflect specialization of the dominant hemisphere or of a particular cerebral hemisphere. Nonetheless, because the available evidence reveals no consistent difference in learning between the two hands of right-handers, it provides no support for either a dominant-hemisphere or a left-hemisphere account, thereby refuting the first prediction of hemispheric specialization accounts of motor learning and instead supporting a functionally symmetric architecture.

### Meta-analysis 2: No Evidence for Asymmetric Interlimb Transfer

We conducted a second meta-analysis designed to address two questions. The first was the extent to which motor adaptation transfers from the trained to the untrained limbs. Answering this question is critical, because interlimb transfer provides a unique window into how motor memories are represented, generalized, and shared across the hemispheres. Second, we asked whether interlimb transfer bears the hallmark signature of hemispheric specialization: greater transfer toward than away from the dominant hand. This prediction follows from the assumption that the dominant hemisphere can learn from errors generated by either hand, whereas the non-dominant hemisphere can only learn from the contralateral hand it more directly controls^51,52,62^.

This meta-analysis yielded 66 datasets derived from 25 studies that met our eligibility criteria (Fig. 2B). Together, these data span over two decades of research (Fig. 3D) and involve 796 participants (Fig. 3F). Following convention, interlimb transfer is defined as the proportion of learning expressed in the untrained limb during the early transfer phase relative to the level of achieved by the trained hand during late adaptation^58^. Under this metric, 0% indicates no transfer, whereas 100% indicates complete transfer of learning between limbs.

The magnitude of interlimb transfer varied substantially across datasets (*τ*^2^ = 0.08, *I*^2^ = 97.7%), ranging between 7.7% and 122.3%. Despite this heterogeneity, the synthesized estimate indicated that approximately 69.7 ± 3.3% of adaptation transferred from the trained to the untrained limb (Fig. 3H; Fig. 5; b= -0.2 ± 0.07, t(21.9) = -3.2, p = 0.004). Thus, although motor adaptation transferred substantially between limbs, it remained incomplete, indicating that motor memories are neither entirely effector-specific nor fully shared across the two limbs.

**Figure 5.**
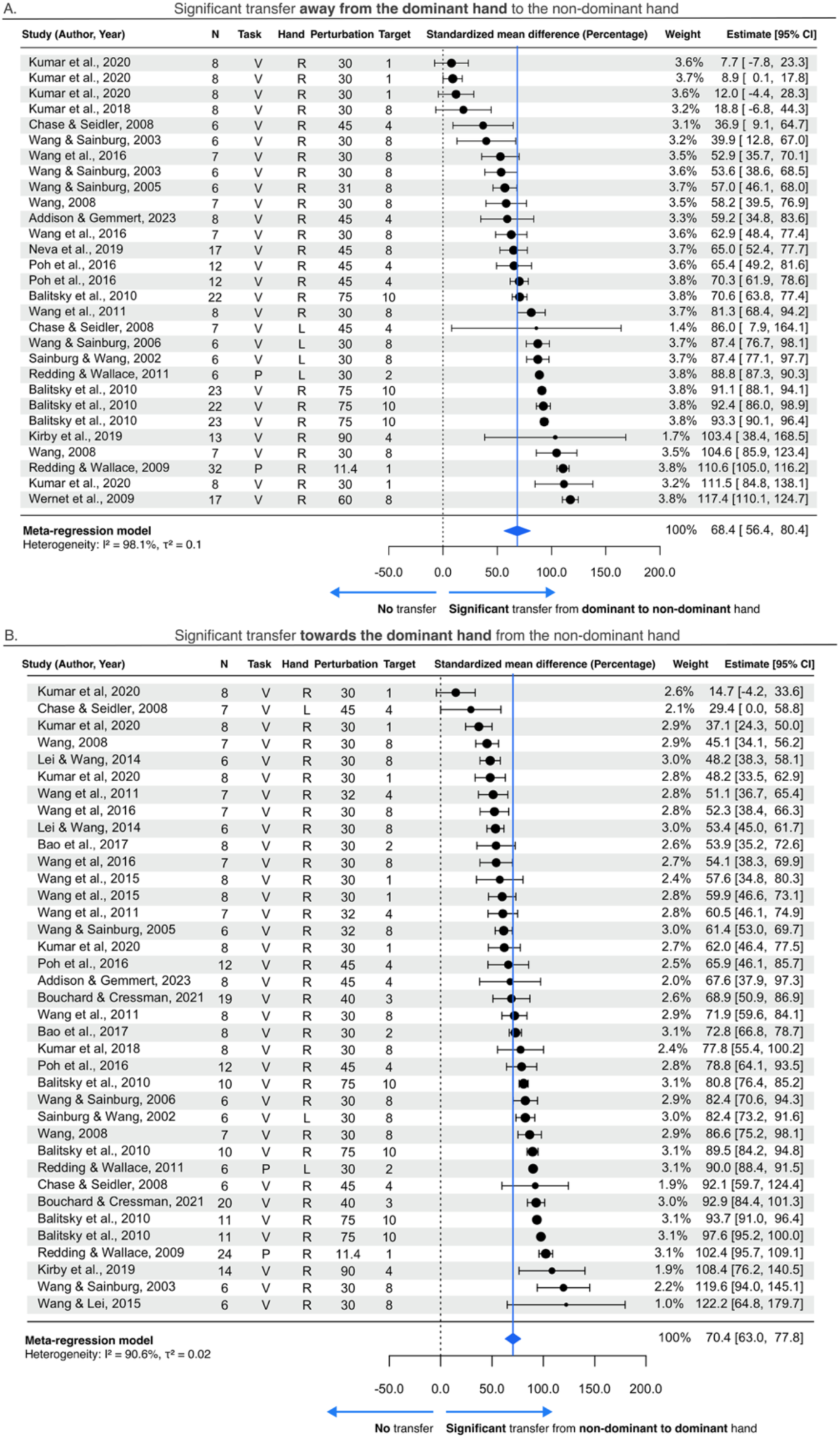
Significant and Symmetric interlimb transfer. **(A-B)** Forest plots aggregate **(A)** 30 datasets quantifying percent transfer from the dominant to the non-dominant limb and **(B)** 38 datasets quantifying transfer from the non-dominant to the dominant limb. The diamond denotes the overall effect size (Hedges’ g) and its 95% confidence interval. The N column shows the number of individuals in the dominant (D) and non-dominant (ND) groups. The size of each effect-size marker reflects the weight assigned to that study in estimating the pooled effect size. In the Task column, V denotes visuomotor adaptation, P prism adaptation, and F force-field adaptation. The Perturbation column indicates the perturbation magnitude (°) used in visuomotor and prism adaptation tasks. The Hand column shows the self-reported handedness of participants. The Target column shows the number of training targets. 0% and 100% denote no transfer and full transfer, respectively.

However, this aggregate estimate likely reflects the combined contribution of multiple learning processes. Evidence for this comes from the only study to cleanly dissociate implicit and explicit learning, in which participants were instructed to either engage or disengage their explicit re-aiming strategy during visuomotor adaptation. Implicit adaptation, measured when participants disengaged their strategy (analogous to an aftereffect), transferred incompletely between limbs (48.3 ± 5.7%), whereas explicit learning, estimated as the difference between total and implicit transfer, transferred completely (105.0 ± 2.9%)^82^. Consequently, the overall estimate of 69.7% transfer in our meta-analysis may represent the average of these distinct forms of learning. More broadly, this finding highlights an important methodological limitation of the existing literature, motivating the need for experiments that isolate implicit and explicit processes to determine their respective contributions to interlimb transfer.

What, then, determines whether learned behavior transfers between limbs? A longstanding hypothesis is that transfer depends on the coordinate frame in which learning is represented. Sensorimotor adaptation is thought to update both extrinsic (visual) and intrinsic (joint-based) representations^83^. Extrinsic representations are accessible to either limb and therefore support interlimb transfer, whereas intrinsic representations are specific to the trained limb and are not readily transferred. Under this framework, implicit adaptation likely transfers incompletely because it reflects motor learning rooted in both extrinsic and intrinsic coordinates^83^. By contrast, explicit strategy use may be rooted predominantly in extrinsic coordinates, and thus, transfers completely.

We examined whether interlimb transfer exhibits the asymmetries predicted by hemispheric specialization. Indeed, there was substantial interlimb transfer in both directions: from the dominant to the non-dominant hand (Fig. 3G & Fig. 5A; synthesized transfer size = 68.4 ± 5.9%; t(14) = -0.9, p = 0.3) and from the non-dominant to the dominant hand (Fig. 3H & Fig. 5B; synthesized transfer size = 70.4 ± 3.7%; t(20.8) = -1.4 ± -0.1, p = 0.2). Importantly, the magnitude of transfer did not differ between directions, providing no evidence for asymmetric interlimb transfer.

Interlimb transfer studies were generally well powered to detect the relatively large transfer effects observed in the literature (Fig. 3F; median statistical power [IQR]: 51.5% [27.9%, 98.5%], with α = 0.05). But none were sufficiently powered to detect asymmetries predicted by hemispheric specialization ([IQR]: 5.4% [5.1%, 7.9%], with α = 0.05). Consequently, the apparent symmetry of interlimb transfer may reflect limited sensitivity rather than the absence of asymmetry. Moreover, the literature focused almost exclusively on right-handed participants and failed to isolate implicit motor learning processes, limitations that we sought to address in our own experiments.

In summary, our meta-analysis revealed that although sensorimotor adaptation transfers from the trained to the untrained limb, there is no evidence of transfer preferentially toward the dominant hand. These findings therefore challenge the second core prediction of the hemispheric specialization accounts of motor learning shown in Figure 1, dominance-advantage and left-advantage, while demonstrating consistency with a functionally symmetric architecture. More direct evidence is provided by the results of the experiments that follow.

### Experiment Overview

Our meta-analysis revealed no evidence for hemisphere-specific advantages in sensorimotor adaptation. However, this conclusion is tempered by two limitations of the existing literature. First, most studies were underpowered, limiting their ability to detect learning asymmetries. Second, the adaptation paradigms used in these studies fail to isolate implicit adaptation—the error-driven process of motor learning, in which the dominant hemisphere is expected to prevail. Because explicit strategies rely on top-down processes that are not intrinsically constrained by the hemisphere controlling the moving limb, they should be acquired equally by either hand and transfer completely to the untrained hand. As a result, explicit motor learning is expected to be relatively symmetric and may therefore obscure putative asymmetries in implicit adaptation.

To overcome these limitations, we conducted a series of preregistered experiments designed to dissociate implicit adaptation from explicit strategy use. Experiment 1 quantified implicit adaptation during discrete goal-directed reaching, whereas Experiment 2 measured explicit strategy use as a point of comparison (combined N = 419) in both left- and right-handers. By testing adaptation across different handedness groups while systematically manipulating the trained hand, these experiments provided a rigorous test of competing theories of the neural organization for motor learning (see Introduction; Fig. 1). Finally, Experiment 3 extended this test to a more dexterous continuous tracking task (N = 107), examining whether the same principles generalize beyond discrete reaching.

Across experiments, we adopted a conservative recruitment strategy, focusing on individuals with strong hand preferences—those scoring at the extremes of the Edinburgh Handedness Inventory (see Methods)^84^. We further verified handedness using objective measures of hand performance, recognizing that self-reported preference does not always correspond to motor proficiency. We first verified that our recruitment strategy identified participants with pronounced motor skill asymmetries in both speeded clicking and goal-directed reaching, consistent with robust dominant-hemisphere lateralization of motor control.

### Dominant-Hemisphere Lateralization in Motor Control

Motor skill was first assessed across hands using a speeded clicking task^85^. Participants rapidly clicked a visual target whose location changed randomly within a grid after each selection (Fig. 6A). Both left- and right-handed participants showed a clear dominant-hand advantage, producing more clicks (Fig. 6B; left-handers: t(401) = 4.6 ± 0.5, p < 0.0001, d = 1.03; right-handers: t(400) = 9.4 ± 0.4, p < 0.0001, d = 2.1) and responding faster with their dominant hand (Fig. 6C; left-handers: t(399) = -96.4 ± 11.2, p < 0.0001, d = -0.9; right-handers: t(400) = -216.8 ± 10.8, p < .0001, d = -2). Consistent with the literature^86–89^, the dominant-hand advantage was more pronounced in right-compared to left-handers (number of clicks, handedness: F(1, 402.95) = 16.2, p < 0.0001; handedness × dominance interaction F(1, 400.18) = 56.4, p < 0.0001; response time, handedness F(1, 403.9) = 18.8, p < 0.0001; handedness × dominance interaction F(1, 399.5) = 59.8, p < .0001).

**Figure 6.**
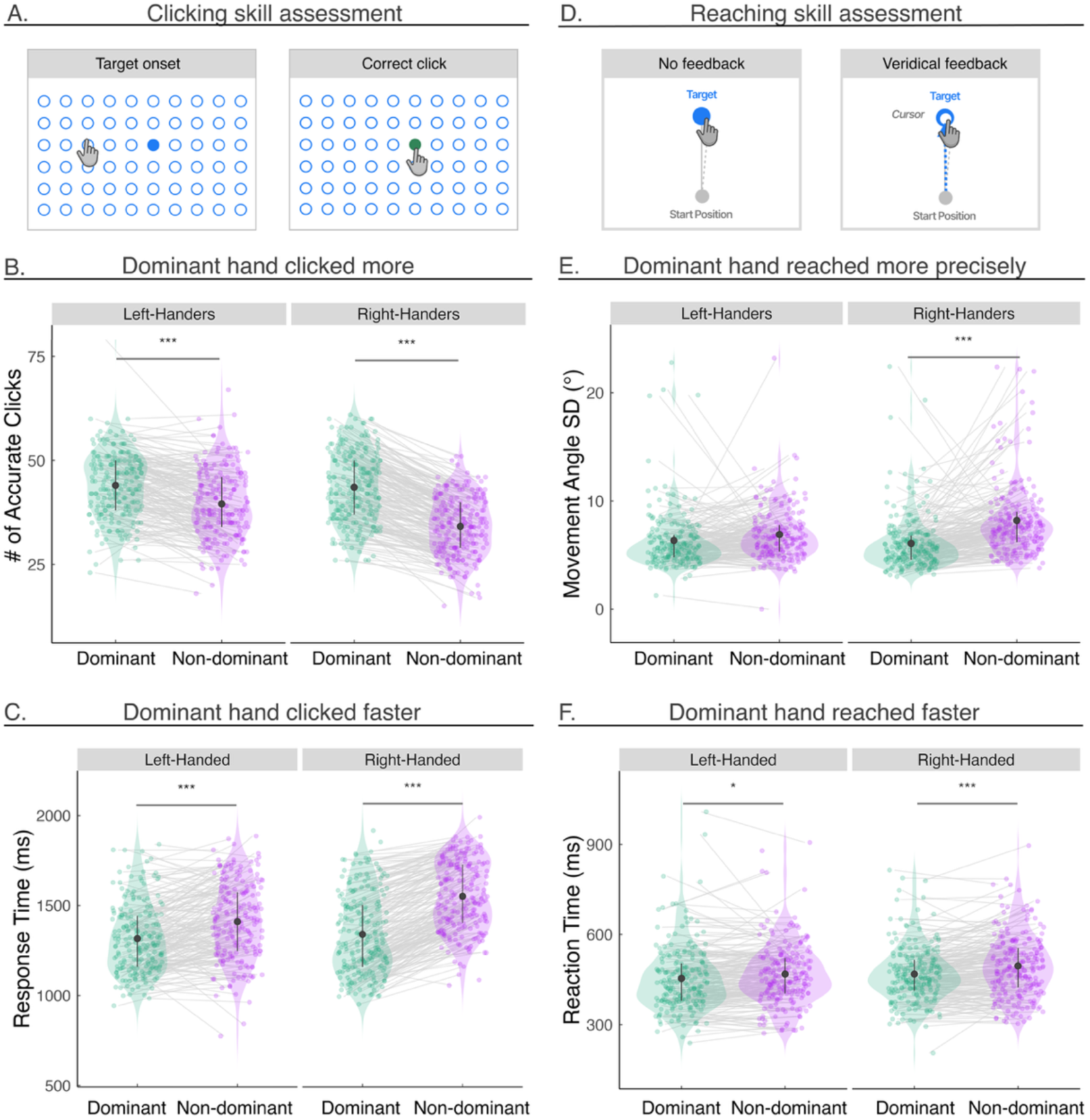
Dominant-Hemisphere Lateralization in Motor Control. **A)** Participants clicked a blue target, randomly selected from a 10 × 6 grid, as quickly and accurately as possible using either their dominant or non-dominant hand. Following a successful click, the target turned green and, after a brief interval, a new blue target appeared. Following an incorrect click, the target turned red before the next target was presented (not shown). **B)** Mean accuracy, quantified by the average number of accurate clicks across two 30-second rounds. **C)** Mean response times, defined as the time from target onset to click. **D)** Schematic of the goal-directed reaching task. Participants were instructed to reach to a visual target with their trackpad or mouse as quickly and accurately as possible, receiving either no visual feedback or veridical feedback, using either their dominant or non-dominant hand. **E)** Mean movement precision, defined as the standard deviation of movement angles. **F)** Mean reaction time, quantified as the time from target onset to movement initiation (crossing 1 cm from the start position). Error bars represent the interquartile range, and the solid dot denotes the group median. Translucent dots represent individual participants. Asterisks denote significant differences between the dominant and non-dominant hands based on paired t-tests (*p < 0.05, **p < 0.01, ***p < 0.001). The width of each violin plot represents the density of the data.

This dominant-hand advantage was further corroborated by a complementary goal-directed reaching task. Participants used a trackpad or mouse to make rapid movements toward visual targets. Consistent with the established literature, right-handers exhibited a clear dominant-hand advantage, reaching both more precisely (Fig. 6E; left-handers: t(396) = -0.5 ± 0.3, p = 0.2, d = -0.2; right-handers: t(388) = -2.1 ± 0.2, p < .0001, d = -0.8) and faster (Fig. 6F; left-handers: t(407) = -12.4 ± 4.6, p = 0.03, d = -0.3; right-handers: t(408) = -26.2 ± 4.4, p < 0.0001, d = -0.6) with their dominant right hand. A similar, albeit trending, dominant-hand advantage was observed in left-handers (accuracy: handedness F(1, 396.5) = 3.4, p = 0.07; handedness × dominance interaction F(1, 392.3) = 19.6, p < 0.0001; reaction time: handedness F(1, 413.9) = 4.5, p = 0.03; handedness × dominance interaction F(1, 407.4) = 4.7, p = 0.03).

In summary, these objective measures confirmed robust dominant-hand advantages across multiple motor control tasks, providing clear evidence for dominant-hemisphere specialization in motor control.

### Experiment 1: Implicit Sensorimotor Adaptation Does Not Exhibit Signatures of Hemispheric Lateralization

In this preregistered experiment, we directly tested whether implicit motor adaptation—the error-driven motor learning process most directly implicated by the dominant-hemisphere hypothesis—is lateralized. To this end, we recruited a large cohort of left- and right-handed participants (Table 1), who were randomly assigned to adapt to a visuomotor perturbation with either their dominant or non-dominant hand. Following adaptation, interlimb transfer was assessed using the untrained hand.

**Table 1.** Demographic Summary. Participants self-identified as Female (F), Male (M) or Prefer Not to Say (N), and as Left-Handed (L) or Right-Handed (R). Reported in the Age column are the average and interquartile ranges per group (age ranges are 18-30 for all experiments). All participants completed the experiments using a computer trackpad.

| Study | Dominant Training Group |  |  |  | Non-dominant Training Group |  |  |  |
| --- | --- | --- | --- | --- | --- | --- | --- | --- |
|  | N | Age | Sex | Handedness | N | Age | Sex | Handedness |
| 1 | 101 | 25.5 ± 0.3<br>[IQR: 23-28] | F = 63,<br>M = 38 | L = 51<br>R = 50 | 101 | 25.4 ± 0.3<br>[IQR: 23-28] | F = 61<br>M = 40 | L = 50<br>R = 51 |
| 2 | 112 | 25.3 ± 0.3<br>[IQR: 22-28] | F = 68<br>M = 44<br>N = 0 | L = 53<br>R = 59 | 105 | 25.4 ± 0.3<br>[IQR: 23-28] | F = 70<br>M = 34<br>N = 1 | L = 50<br>R = 55 |
| 3 | 53 | 25.4 ± 0.5<br>[IQR: 23-28] | F = 23<br>M = 29<br>N = 1 | L = 0<br>R = 53 | 54 | 25.5 ± 0.5<br>[IQR: 23-28.8] | F = 23<br>M = 30<br>N = 1 | L = 0<br>R = 54 |

To isolate implicit adaptation, we employed a clamped feedback task in which the visual cursor followed an invariant trajectory on each trial: its radial position was yoked to the participant’s movement, while its angular position was fixed at a constant offset relative to the target (Fig. 7A). Participants were instructed to ignore this non-contingent feedback and reach straight (“slice”) through the target. Despite these instructions, movement angles in all groups gradually shifted in the direction opposite the cursor rotation—a signature of implicit adaptation. Moreover, the magnitude of late adaptation closely matched the no-feedback aftereffect phase, providing further evidence that this task isolates this implicit learning process^90–92^.

**Figure 7.**
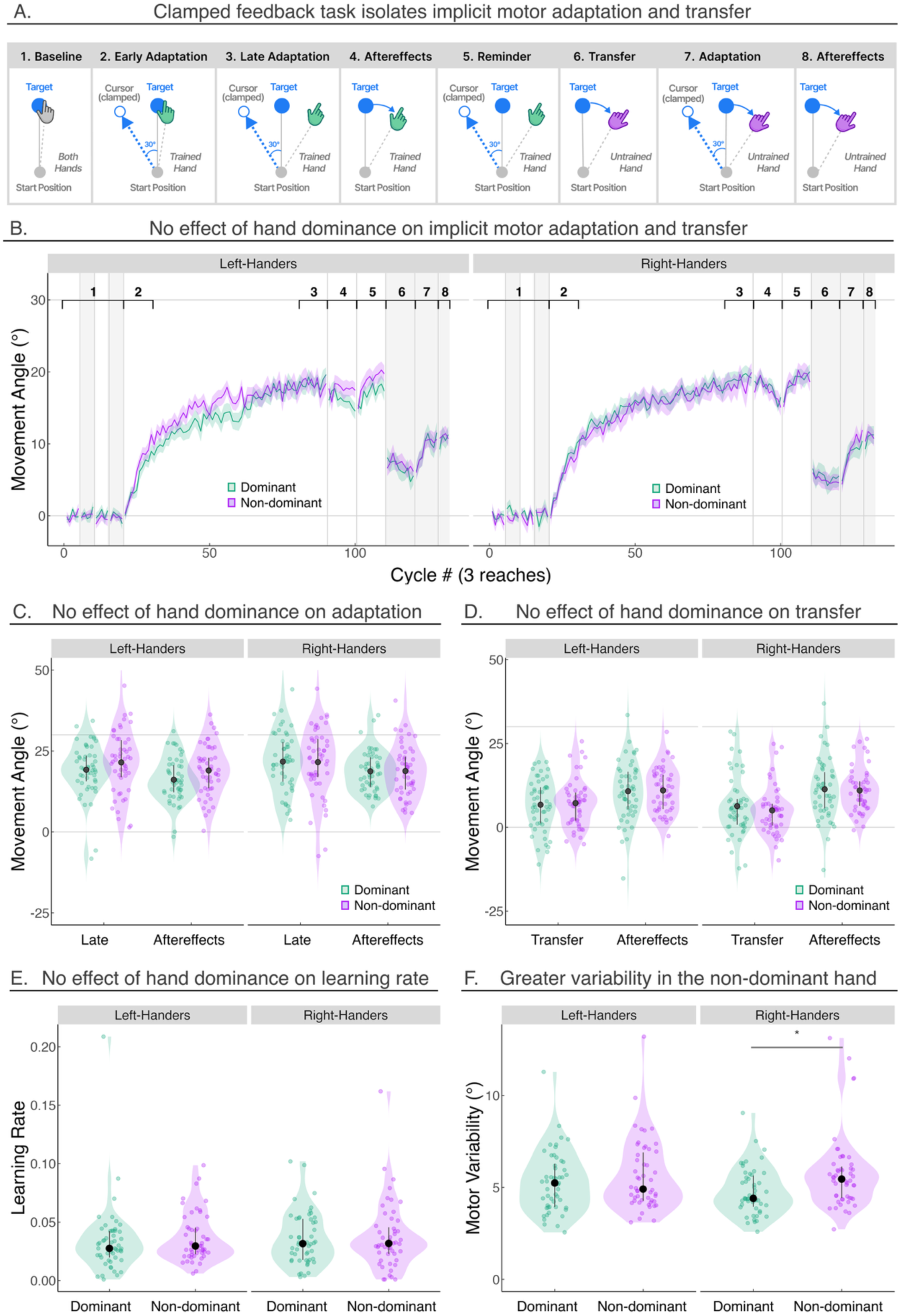
Implicit Sensorimotor Adaptation Does Not Exhibit Hemispheric Lateralization. **A)** Schematic of the clamped feedback task used to isolate implicit adaptation (Experiment 1). Participants were randomly assigned to adapt to a visuomotor perturbation with either their dominant or non-dominant hand and subsequently switched to their untrained hand to assess interlimb transfer. Participants were always asked to always reach straight to the target (filled blue dot). During the clamped feedback phase, a cursor rotated by 30° (hollow blue circle) was displayed throughout the movement. Participants were instructed to ignore this feedback; nevertheless, their movement angles gradually deviated away from the target, the signature of implicit adaptation. **B)** Mean learning curves for participants using the dominant or non-dominant hand across task phases (black numbers). The purple curve denotes the group trained with the non-dominant hand, whereas the green curve denotes the group trained with the dominant hand. Light gray bars indicate blocks where the participants assigned to the dominant and non-dominant hand training group used their other, untrained hand. **C-D)** Mean movement angles during initial adaptation phases (phases 3 and 4) and interlimb transfer phases (phases 6 and 8). **E-F)** Learning rate and motor noise derived from fitting the state space model. Error bars represent the interquartile range, and the solid dot denotes the group mean. Translucent dots represent individual participants. Stars denote significant differences (*p < 0.05, **p < 0.01, ***p < 0.001) based on t-tests between dominant and non-dominant hands. The width of each violin plot represents the density of the data.

Statistical analyses confirmed that all four groups exhibited a gradual shift in movement angle away from the target, hitting an asymptote around 15° - 25°, a range consistent with prior literature^93^ (Fig. 7B; left dominant t(198) = 19.1 ± 1.3, p < 0.0001, d = 3; left non-dominant t(198) = 21.7 ± 1.3, p < 0.0001, d = 3.4; right dominant t(198) = 21.6 ± 1.3, p < 0.0001, d = 3.2; right non-dominant t(198) = 22 ± 1.3, p < 0.0001, d = 3.5). For all groups, performance during the aftereffect phase showed only a modest decrement relative to late adaptation (left dominant t(198) = -3.1 ± 0.7, p < 0.0001, d = -0.8; left non-dominant t(198) = 2.5 ± 0.7, p = 0.0008, d = -0.7; right dominant t(198) = -2.9 ± 0.7, p = 0.000q, d = -0.8; right non-dominant t(198) = -2.7 ± 0.7, p = 0.0002, d = -0.7).

We next addressed our primary question: does implicit adaptation exhibit a dominant-hand advantage? Notably, the inclusion of both left- and right-handed participants provided a particularly stringent test of the dominant-hemisphere hypothesis. If motor adaptation mirrors motor control, learning advantages should track dominant-hand superiority: right-handed individuals should exhibit greater adaptation with their right hand, whereas left-handed individuals should exhibit greater adaptation with their left hand. Moreover, the magnitude of this learning advantage should scale with the degree of motor lateralization, predicting a smaller dominant-hand advantage in left-handed participants, consistent with their weaker asymmetries in motor skill (Fig. 6).

Across multiple analyses, we found no evidence for a dominant-hand advantage in either left- or right-handers, nor did we find indications of a general left-hand learning deficit as predicted by the left-hemisphere advantage account in Figure 1b. First, using conventional linear modeling, neither handedness nor hand dominance significantly affected late adaptation (Fig. 7B, C; handedness F(1, 196) = 0.9, p = 0.3, BF = 0.2 ± 0.1% in favor of the null; hand dominance F(1, 196) = 0.7, p = 0.4, BF = 0.2 ± 0.1% in favor of the null). Post hoc comparisons likewise revealed no dominant-hand advantage in either group (left-handers: t(195) = -2.1 ± 1.9, p = 0.7, d = -0.3; right-handers t(195) = -0.2 ± 1.9, p = 1.0, d = -0.02). Controlling for individual differences in motor performance (reaction time and movement time) did not alter the null effects of handedness and hand dominance (F(1, 195) = 0.3, p = 0.6, BF = 0.02 ± 1.8%).

Second, a more sensitive cluster-based permutation analysis comparing dominant and non-dominant hands across the entire adaptation time course revealed no significant clusters, indicating that learning trajectories were statistically indistinguishable between hands. Third, a model-based analysis reached the same conclusion (see Methods): estimated learning rates did not differ as a function of handedness (Fig. 6E; F(1, 198) = 0.1, p = 0.7, BF = 0.2 ± 0.1%) or hand dominance (F(1, 198) = 0.5, p = 0.5, BF = 0.2 ± 1.5%). Together, these converging analyses provide no evidence for a dominant advantage in implicit adaptation.

Importantly, these null effects in implicit adaptation were observed despite robust dominant-hand advantages in motor control. Right-handed participants exhibited significantly greater motor variability when using their nondominant hand during baseline reaching (Fig. 6), a pattern that was independently confirmed by our model-based estimates of motor variability (Fig. 7F; t(198) = -0.9 ± 0.3, p = 0.03, d = -0.6). Together, these findings reveal a striking dissociation between motor control and motor learning: whereas the control of skilled action is strongly specialized to the dominant hemisphere, the capacity for motor learning appears to function symmetrically across the two hemispheres.

Having established that implicit learning is equated whether the trained hand is dominant or nondominant, and across both right- and left-handed individuals, we next asked whether implicit adaptation transfers asymmetrically between the limbs. After the initial learning phase, participants were asked to switch to their untrained hand and, without visual feedback, instructed to continue reaching to the target without engaging any re-aiming strategy. If adaptation transfers across limbs, an aftereffect should be observed in the untrained hand. Moreover, if the neural correlates of hand dominance described earlier – greater excitability, efficiency, and connectivity of the dominant hemisphere – lead to an advantage in implicit motor learning, transfer should be greater transfer *toward* versus *away* from the dominant hand.

We observed significant, yet partial, interlimb transfer, averaging 40.8 ± 5% across participants (Fig. 7D). Notably, this estimate of the percentage of learning transferred was substantially lower than that observed in our meta-analysis (69.6% ± 3.3%), likely reflecting the fact that most studies in the literature do not isolate implicit learning processes. Our results thus provide an uncontaminated estimate of implicit interlimb transfer, verifying that implicit motor memories comprise both intrinsic (joint-based) and shared extrinsic (visual) representations^83^.

Critically, interlimb transfer did not vary as a function of transfer direction (F(1, 198) = 0.01, p = 0.9, BF = 0.2 ± 0.07%) or handedness (Fig. 7D; F(1, 53.3) = 0.8, p = 0.4, BF = 0.2 ± 0.05%): On average, transfer from the dominant to the non-dominant hand was 39.4 ± 5.6%, and transfer from the non-dominant to the dominant hand was 42.2 ± 8.3%, indicating highly similar levels of transfer in both directions. Implicit adaptation thus transferred symmetrically between the limbs. Taken together, the absence of both a dominant-hand advantage for initial training and asymmetric interlimb transfer argues against a hemispheric-specialization advantage for implicit motor learning (Fig. 1A, B). We will consider the neural basis of this functional symmetry in the Discussion section.

### Experiment 2: Explicit Strategy Use Does Not Exhibit Signatures of Hemispheric Lateralization

Although theories of hemispheric specialization have focused almost exclusively on implicit adaptation, whether explicit strategy use exhibits comparable hemispheric asymmetries has remained largely unexplored. Unlike implicit adaptation, which has been hypothesized to rely on effector-specific sensorimotor processes, explicit strategy use depends on higher-order cognitive representations that can, in principle, be acquired by and transferred to either hand. Accordingly, we predicted that explicit strategy use would exhibit little or no behavioral asymmetry, consistent with a functionally symmetric neural organization.

To test this, we recruited a large cohort of left-(N = 103) and right-handed participants (N = 114), who adapted to a 60° visuomotor rotation with either their dominant or nondominant hand before interlimb transfer was assessed in the previously untrained hand. To isolate strategy use, we employed a classic delayed-feedback paradigm in which endpoint feedback was presented 800 ms after movement completion. This manipulation minimizes implicit adaptation, thereby biasing learning toward explicit re-aiming strategies (Fig. 8A)^90,92,94,95^.

**Figure 8.**
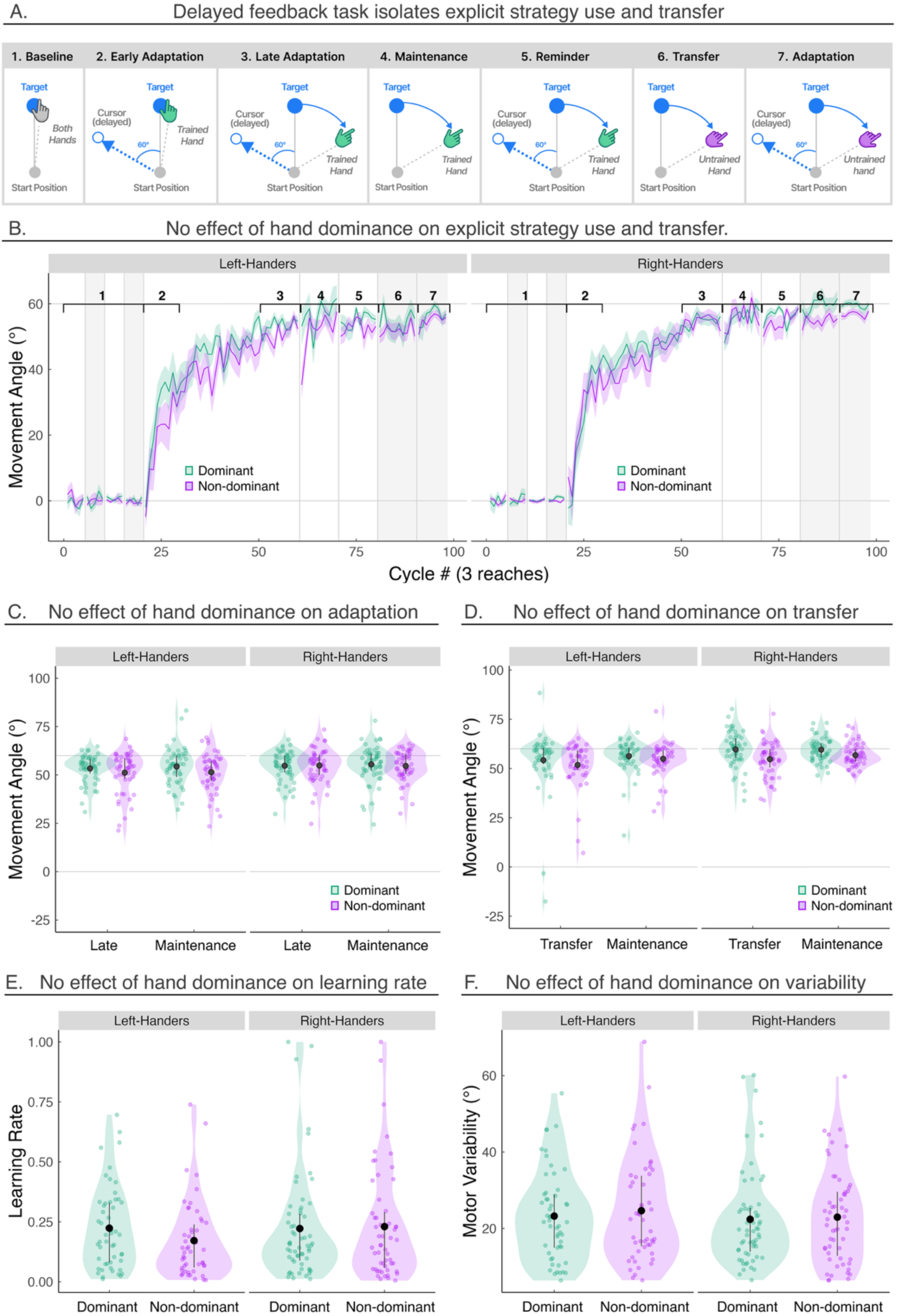
Explicit Strategy Use Does Not Exhibit Signatures of Hemispheric Lateralization. **A)** Schematic of the delayed feedback task used to assess explicit motor adaptation (Experiment 2). Participants were randomly assigned to train with either their dominant or non-dominant hand and were subsequently asked to express what they had learned using the previously untrained hand to assess interlimb transfer. Throughout the experiment, participants were instructed to select a movement such that the visual cursor “hit” the target. During the delayed-feedback phase, visual feedback consisted of a cursor rotated 60° relative to the target (blue circle), presented at the movement endpoint 800 ms after participants reached the target distance. **B)** Mean learning curves for participants using the dominant or non-dominant hand across task phases (black numbers). The purple curve denotes the group trained with the non-dominant hand, whereas the green curve denotes the group trained with the dominant hand. **C-D)** Mean movement angles during adaptation phases (phases 3 and 4) and transfer phases (phases 6 and 7). Error bars represent the interquartile range, and the solid dot denotes the group mean. Translucent dots represent individual participants. For clarity, data from three outlier participants whose movement angles exceeded 100° during the late adaptation and maintenance phases are not shown but were included in all calculations of means and standard errors. **E-F)** Learning rate and motor noise derived from fitting the state space model. Error bars represent the interquartile range, and the solid dot denotes the group median. Translucent dots represent individual participants. Stars denote significant differences (*p < 0.05, **p < 0.01, ***p < 0.001) based on t-tests between dominant and non-dominant hands. The width of each violin plot represents the density of the data.

Participants in all groups successfully discovered and implemented re-aiming strategies to compensate for the visuomotor rotation, reaching late adaptation levels of approximately 50°–55° (Fig. 8B; left dominant t(213) = 53.2 ± 1.2, p < 0.0001, d = 8.4, left non-dominant t(213) = 50.7 ± 1.3, p < 0.0001, d = 7.9, right dominant hand t(213) = 54.8 ± 1.2, p < 0.0001, d = 8.6, and non-dominant hand t(213) = 54.8 ± 1.2, p < 0.0001, d = 8.6). Importantly, our previous work using this same paradigm revealed negligible aftereffects, confirming that learning under these conditions is overwhelmingly driven by explicit processes^96,97^. Consequently, to minimize task duration given the multiple phases of the protocol, we did not include a no-feedback aftereffect phase.

We next asked whether explicit strategy use exhibits the signatures of hemispheric specialization. Overall, participants successfully implemented explicit re-aiming strategies to counter the visuomotor rotation. Although there was a statistically significant effect of handedness (Fig. 8C; F(1, 210) = 6.3, p = 0.01, BF = 1.4 ± 0.01%), the magnitude of this difference was small, with left-handers exhibiting only 2.8 ± 1.1° less adaptation than right-handers (t(213) = -2.5 ± 1.2, p = 0.03, d = -0.3; significant group differences identified during cycles 80-81, 85-88, and 90-94). But this effect was not evident in model-based estimates of learning rate (F(1, 0.04) = 1.1, p = 0.3, BF = 4.4 ± 1.1%), suggesting that any differences between left- and right-handed participants were negligible.

Critically, we found no evidence for the dominant-hand advantage in strategy use. Late adaptation did not differ as a function of hand dominance (F(1, 213) = 0.8, p = 0.4, BF = 0.2 ± 0.05%), either in left-handers (t(213) = 2.2 ± 1.7, p =0.6, d = 0.3) or right-handers (t(213) = -0.1 ± 1.6, p = 0.9, d = -0.02). The absence of a dominant-hand advantage persisted during the subsequent strategy maintenance phase (F(1, 213) = 2.3, p = 0.1, BF = 0.4 ± 0.04%), indicating that neither the discovery nor short-term retention of explicit strategies was preferentially expressed in the dominant hand.

This null effect of hand dominance was reinforced by cluster-based permutation analyses, which revealed no significant differences between dominant and non-dominant hands at any point during adaptation. Consistent with this result, model-based estimates of learning rate showed no effect of hand dominance (F(1, 213) = 0.03, p = 0.9, BF = 0.2 ± 0.%). Together, these converging analyses provide no evidence for the dominant-hand advantage predicted by hemispheric specialization accounts.

We next examined the extent to which strategy use transfers between the limbs and, critically, whether such transfer exhibits the asymmetries predicted by hemispheric specialization. When participants were asked to express their strategy using their untrained limb, participants exhibited complete transfer of explicit strategies, averaging 104.2 ± 1.9% (Fig. 8D). This complete transfer is consistent with the idea that strategic learning operates at an abstract, extrinsic, and effector-independent level, with little contribution from effector-dependent, intrinsic, or joint-based representations^82,98^.

Importantly, the magnitude of interlimb transfer did not vary as a function of direction (Fig. 8D; F(1, 213) = 0.001, p = 0.9, BF = 0.1 ± 0.08%) and was equal to 104.1 ± 1.9%. Transfer from the dominant to the non-dominant hand averaged 106.9 ± 2.4%, whereas transfer from the non-dominant to the dominant hand averaged 101.2 ± 2.8%, with largely overlapping distributions across groups. This effect is not due to a ceiling effect, as substantial inter-individual variability was evident within each group. Thus, explicit strategies transferred symmetrically between the limbs, providing no evidence for the asymmetric interlimb transfer.

In summary, the absence of both a dominant-hand advantage and asymmetric interlimb transfer argues against both dominant-hemisphere and left-hemisphere accounts for explicit motor learning. We will consider the neural basis of this functional symmetry in the Discussion.

### Experiment 3: Sensorimotor Adaptation During Continuous Tracking Does Not Exhibit Signatures of Hemispheric Lateralization

The absence of hemispheric specialization observed in Experiments 1–2 may reflect the relatively simple nature of the discrete reaching tasks, which may not sufficiently engage the computational demands that give rise to lateralized motor control. If hemispheric specialization represents a general organizing principle of motor learning, it may be more likely to emerge during dexterous, continuous, and naturalistic behaviors that more closely resemble the complexity of everyday motor learning^99^.

To test this, we employed a more naturalistic adaptation paradigm in which right-handed participants used either their dominant or non-dominant hand to track a continuously moving target for 40 seconds per trial (N = 107, Table 1). We did not recruit left-handed participants because our preceding experiments revealed no evidence for handedness effects on learning. Following a baseline phase, a visuomotor rotation was gradually introduced (+1° per trial) to a maximum of 30° (Fig. 9A). Because gradual perturbations minimize awareness and strategy use, adaptation is thought to be driven predominantly by implicit learning processes^100^. Moreover, continuous tracking engages both feedforward and feedback processes, providing a rich and ecologically valid assay of motor learning that more closely reflects the multiple control processes underlying real-world behavior.

**Figure 9.**
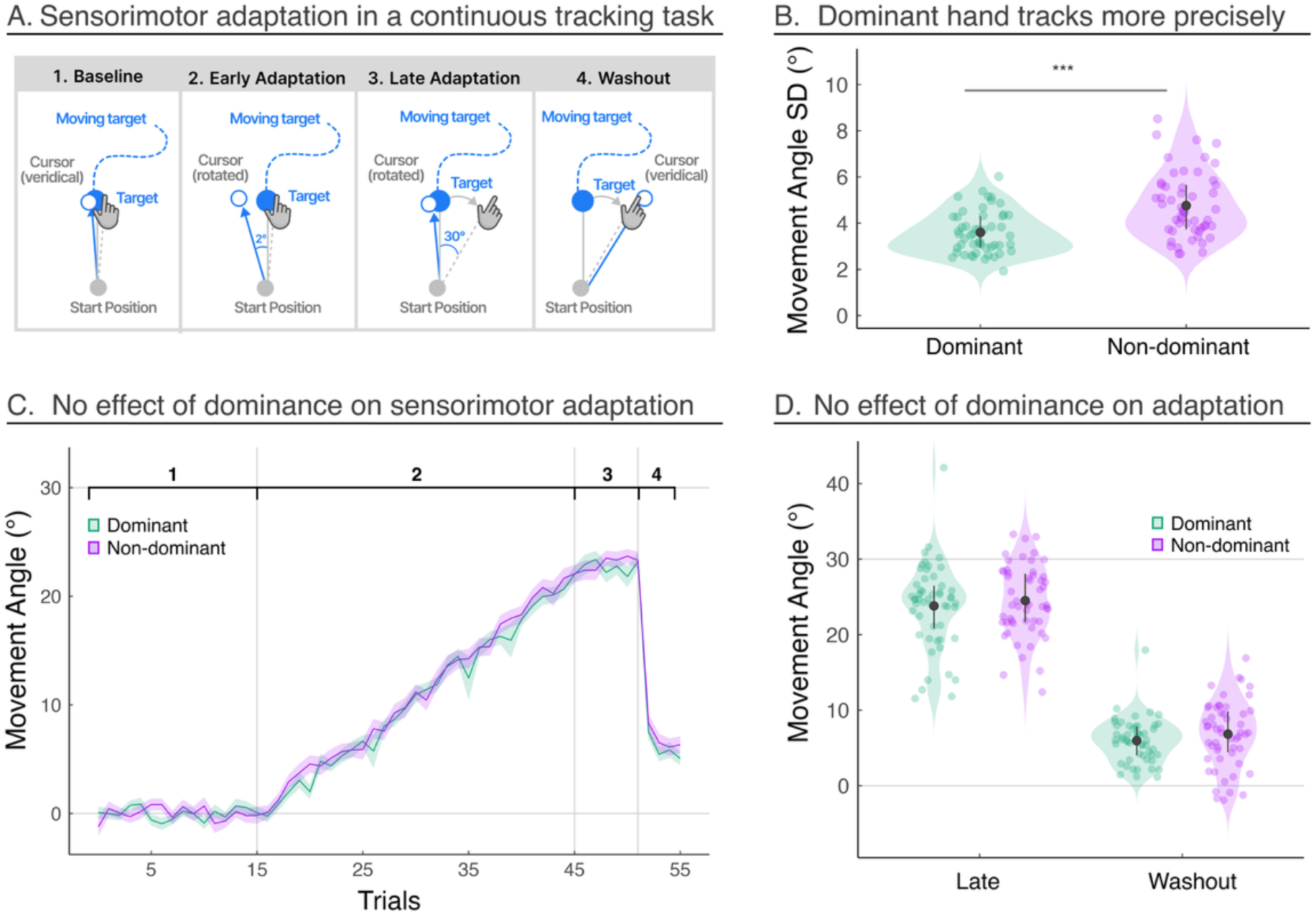
Sensorimotor Adaptation During Continuous Tracking Does Not Exhibit Signatures of Hemispheric Lateralization. **A)** Schematic of the continuous tracking task. Participants were randomly assigned to perform the task with either their dominant or non-dominant hand. They were instructed to continuously track a target moving along a sum-of-sines trajectory. During early adaptation, a perturbed cursor (hollow blue circle) was displayed throughout the movement, with the visuomotor rotation gradually increasing across trials to 30°. During late adaptation, the perturbation remained fixed at 30°. During the aftereffect phase, cursor feedback returned to veridical, enabling measurement of learning-related aftereffects while preserving continuous task engagement. **B)** Movement variability, quantified as the standard deviation of movement angles, during the baseline phase with veridical feedback for dominant- and non-dominant-hand groups. **C)** Mean learning curves for participants using the dominant (green) or non-dominant hand (purple) across task phases (black numbers). **D)** Mean movement angles during late adaptation and aftereffects phases (phases 3 and 4). Error bars represent the interquartile range, and the solid dot denotes the group mean. Translucent dots represent individual participants. Stars denote significant differences (*p < 0.05, **p < 0.01, ***p < 0.001) based on t-tests between dominant and non-dominant hands. The width of each violin plot represents the density of the data.

We first examined performance during the baseline phase to determine whether motor control was lateralized. Consistent with this possibility, the non-dominant hand displayed significantly greater motor variability than the dominant hand (t(102) = -1.2 ± 0.2, p < 0.0001, d = -0.9), indicating reduced movement precision (Fig. 9B). This difference could not be explained by a speed–accuracy trade-off, as average movement velocity was comparable across hands (t(141) = -38.3 ± 26.4, p = 0.1, d = -0.6). Thus, these findings demonstrate a robust dominant-hand advantage in motor execution, providing a positive control that the task was sensitive to well-established forms of hemispheric specialization.

We next asked whether sensorimotor adaptation during continuous tracking exhibited a similar dominant-hand advantage. Strikingly, it did not. Adaptation to the gradual perturbation was nearly identical across hands throughout the learning phase (no significant clusters found). During the late phase, participants compensated for the perturbation to a similar degree with their dominant (23.8° ± 0.8°) and non-dominant (24.5° ± 0.6) hands (Fig. 9C-D; t(383) = -0.7 ± 0.7, p = 0.3, d = -0.2). Likewise, both groups exhibited comparable levels of aftereffect during the washout phase (Fig. 9C-D; dominant: 5.9° ± 0.4°, non-dominant: 6.8° ± 0.6°; t(85) = -0.9 ± 0.7, p = 0.2, d = -0.3), indicating similar degrees of implicit adaptation. Together, these findings provide further evidence against a dominant-hand advantage in sensorimotor adaptation, consistent with a functionally symmetric neural architecture for motor learning.

## Discussion

Many core human functions are organized asymmetrically across the two hemispheres, with one playing a dominant role. This principle is perhaps most evident in the motor system, where motor control is strongly lateralized, giving rise to well-established dominant-hand advantages in movement execution^1,3,4,14^. Efforts to readily extend patterns of dominant advantages in motor control to motor learning have not been successful^64,65,67,101^. Here, we revisited hemispheric organization of motor learning through complementary meta-analyses and preregistered experiments that isolated implicit and explicit sensorimotor adaptation in a large, well-characterized cohort of strongly left- and right-handed individuals across both discrete and continuous motor tasks.

Across complementary meta-analytic and experimental approaches, the evidence consistently contradicted the core predictions of dominant-hand advantages for motor learning. Right- and left-handed participants adapted at the same rate and to the same extent regardless of which hand was trained, and learning transferred symmetrically between hands across discrete and continuous adaptation contexts. Importantly, this functional symmetry does not appear to be unique to visuomotor adaptation. Previous work has shown that force-field adaptation is likewise acquired symmetrically across the two hands, indicating that this principle extends beyond updating visuomotor mappings to the learning of intersegmental limb dynamics^102^. Nor is functional symmetry restricted to discrete movements: here, we observed equivalent adaptation during both ballistic reaching and dexterous target tracking tasks. Nor is functional symmetry limited to sensorimotor adaptation. Converging evidence from motor skill learning likewise demonstrates equivalent improvements of the dominant and non-dominant hands when initial performance is matched^64,103–107^. Together, these findings suggest that functional symmetry is not a task-specific phenomenon, but rather a general organizing principle of motor learning.

Importantly, our claim is one of *functional* symmetry: Our results indicate that the neural representations supporting explicit strategy use and implicit motor adaptation are updated equally regardless of which hand is trained. Functional symmetry does not dictate neural architecture but rather constrains it. One possibility is that functional symmetry arises from a neural architecture that does not exhibit lateral dominance. This might be a central resource that networks across the two hemispheres or separate centers in each hemisphere that perform equivalently (our working hypothesis; Fig. 10). An alternative, albeit less parsimonious, possibility is that the underlying sensorimotor computations are lateralized, but their outputs are equally accessible to both hands. We next consider these possibilities in more detail.

**Figure 10.**
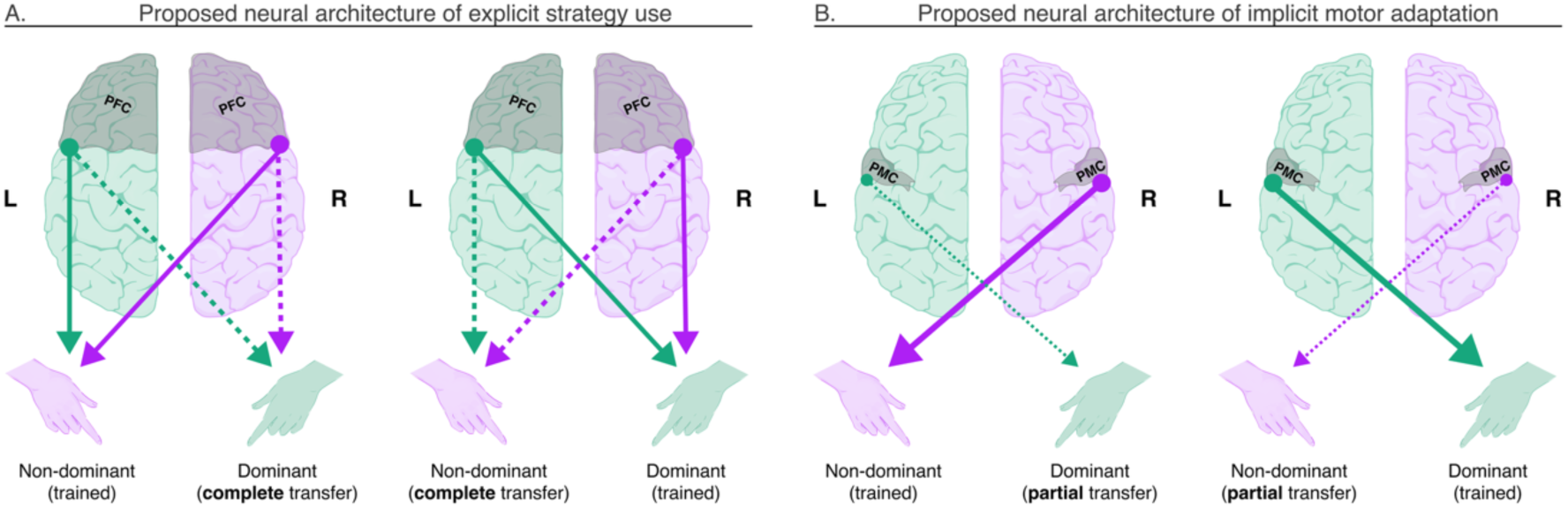
Proposed symmetric neural architecture for motor learning (right-handers). **(A)** Strategy use during motor learning is learned symmetrically between hands and transfers completely and symmetrically between hands, supported by the prefrontal cortex (PFC). **(B)** Implicit motor learning is learned symmetrically between hands and transfers partially, but symmetrically, between hands, supported by the premotor cortex (PMC). Solid arrows denote the proposed contributions of each region to motor learning, whereas dashed arrows denote proposed contributions to interlimb transfer. Arrow thickness represents the relative magnitude of learning and transfer.

### What neural architecture(s) could give rise to the functional symmetry observed in motor learning?

We first consider explicit strategy use. Converging evidence has long implicated the prefrontal cortex (PFC) in strategic motor learning (Fig. 10A). Neuroimaging studies consistently demonstrate robust PFC engagement during strategy use^108^, whereas patients with PFC lesions exhibit selective strategic impairments while largely sparing implicit adaptation^109,110^. Importantly, the PFC exhibits largely symmetric connectivity with bilateral sensorimotor regions^111^, providing a plausible substrate for effector-independent representations that, once acquired, are equally accessible to either hand^83,112–114^. Such an architecture naturally explains the symmetric acquisition and transfer of explicit strategies observed here.

We next consider implicit motor learning. Converging evidence likewise implicates premotor cortex (PMC) as a key substrate for implicit sensorimotor adaptation (Fig. 10B). Neuroimaging studies consistently demonstrate robust PMC engagement during implicit adaptation^59,115,116^, while causal evidence from noninvasive brain stimulation shows that disrupting PMC selectively impairs implicit adaptation^117–119^. Moreover, implicit adaptation is thought to update sensorimotor mappings in both effector-independent (visual-based) and effector-dependent (joint-based) coordinate frames^82,113,114^. Consequently, adaptation gives rise to both shared and limb-specific sensorimotor representations. The shared component provides a natural explanation for the substantial, symmetric interlimb transfer observed here, whereas the limb-specific component explains why transfer remains incomplete.

While prefrontal cortex (PFC) and premotor cortex (PMC) can be identified as plausible neural substrates supporting strategy use and implicit adaptation, respectively, identifying a neural locus does not resolve the issue of the nature of the representation that produces a functional symmetry. As noted previously, the symmetric pattern of learning could arise from an architecture that does not exhibit lateral dominance, in the form of a central resource or bilateral structures that perform equivalently^120–123^ (our working hypothesis; Fig. 10). Either of these hypotheses could be applied to PFC or PMC, although it is an intriguing possibility that PFC might function centrally^124–126^. and PMC functions might be duplicated laterally^123,127^. A less parsimonious alternative is a lateralized architecture whose representations are equally accessible to both hands, which again might apply to PFC or PMC^128^. Our behavioral findings cannot distinguish between these candidate architectures. Resolving this question will require future causal experiments that selectively perturb each hemisphere during motor learning.

In contrast to the symmetry of motor learning, motor control was strikingly asymmetric. Across both discrete and continuous tasks, participants executed speeded, goal-directed movements faster and more accurately with their dominant hand, consistent with the well-established lateralization of motor execution to the dominant hemisphere. This behavioral asymmetry may arise due to a specialized dominant primary motor cortex that has denser structural and functional connectivity within the dominant sensorimotor network as well as larger cortical representations of and stronger corticospinal projections to the contralateral dominant limb^3,76,129–131^.

One particularly striking aspect of our findings is that implicit motor learning was equivalent across the two hands, despite the non-dominant hand exhibiting substantially greater movement variability during baseline. This was especially evident in Experiment 3, where continuous tracking with the non-dominant hand produced markedly noisier movement trajectories, yet implicit adaptation remained equally robust. This resilience is consistent with recent proposals that implicit adaptation operates on an internal estimate of motor output that compensates for execution noise^132–135^, highlighting the remarkable ability of the sensorimotor system to isolate learning-relevant error signals from variability generated by its own motor commands.

### Yet this dissociation between motor control and motor learning raises an apparent paradox: how can pronounced asymmetries in motor control emerge from functionally symmetric motor learning?

Our findings argue against the strong hypothesis that dominant-hand advantages arise *because* the dominant hemisphere is innately equipped with a superior capacity for motor learning, which in turn gives rise to superior motor control. One possibility is that hemispheric specialization in motor control is instead established through innate mechanisms that are largely independent of motor learning. Another intriguing proposal is that there are no intrinsic differences between the hemispheres in motor control or learning capacity, but instead, asymmetries in skilled motor behavior emerge primarily through differential feedback-governed experience^103^. By this account, small biological predispositions initially bias hand selection for everyday tasks such as writing, eating, and tool use, and these predispositions act over years of asymmetric experience to ultimately generate large differences in motor control despite functionally symmetric mechanisms in motor learning.

Taken together, these findings refute the long-standing assumption that dominant-hand advantages in motor control necessarily reflect a lateralized advantage in motor learning. Instead, they suggest that motor control and motor learning follow fundamentally different organizational principles: Whereas motor execution is lateralized to the contralateral hemisphere, motor learning appears to operate through neural architecture that supports equivalent learning and transfer across the two hands.

## Methods

### Code and data availability

All code and data are shared publicly as part of the OpenMotor Database (https://osf.io/aknqj).

### Meta-analysis

#### Study identification, screening, and eligibility

We conducted two meta-analyses: the first tested whether motor learning is preferentially expressed in the dominant hand, whereas the second examined whether interlimb transfer is asymmetric, favoring transfer toward rather than away from the dominant limb. At each stage, we adhered to the Preferred Reporting Items for Systematic Reviews and Meta-Analyses (PRISMA) guidelines. We conducted systematic literature searches in PsycINFO, PubMed, ProQuest, and Google Scholar, and identified additional peer-reviewed publications through social media, conference proceedings, and recommendations from colleagues.

For both meta-analyses, we searched for studies between 1900 and 2025 using search terms in Supplementary Materials. This search yielded 1,915 records for the learning-focused meta-analysis and 892 records for the transfer-focused meta-analysis. Three authors (IN, AS, EM) independently screened records, removed duplicates, and assessed eligibility. Published studies were included if they (1) measured upper-limb adaptation in response to a visual perturbation (e.g., visuomotor rotation or prism adaptation) or a force perturbation, (2) involved neurologically healthy participants, and (3) were written in English; (4) For the first meta-analysis, studies were required to compare adaptation between the dominant and non-dominant limbs; for second meta-analysis, studies were required to include a measure of transfer to the untrained limb. The screening results for each analysis are summarized in Figure 1.

#### Motor adaptation tasks

Our meta-analysis included two types of motor adaptation tasks that differed in the nature of the perturbation. In visual perturbation tasks (e.g., visuomotor rotation or prism adaptation), participants reach to a visual target and receive visual feedback that is perturbed (e.g., rotated or translated) with respect to the position of the hand (Fig. 2A). Over the course of learning, participants nullify this perturbation by moving the limb in the opposite direction of the perturbation, drawing the perturbed visual feedback closer to the target. In a subsequent no-feedback post-learning block, the perturbation and feedback are removed, and participants are instructed to abandon any explicit strategies. Despite these instructions, participants often exhibit a persistent change in movement angle in the direction of learning, also known as the “aftereffect”.

In force-field perturbation tasks, participants make a goal-directed reach to a visual target while holding a robotic device with feedback again restricted to a cursor indicating the participant’s hand position (Fig. 2A). During the perturbation phase, the robot arm applies a velocity-dependent or position-dependent force that displaces the participant’s hand away from the target, resulting in a curved hand trajectory. Over the course of learning, participants learn to exert an equal and opposite force to counteract the force-field perturbation, with the end result being movement trajectories that are relatively straight from the start position to the target. During a subsequent post-learning block, the perturbation and feedback are removed and participants exhibit an aftereffect, now expressed as moments displaced to the opposite side of that observed in the early phases of adaptation.

In both task types, participants may be asked to transfer what they learned with the trained limb to the untrained limb (Fig. 2B). However, there is substantial variability in how this transfer phase is implemented. Specifically, studies differ in the instructions provided: some simply ask participants to continue performing the task with the opposite limb, leaving it ambiguous whether explicit re-aiming strategies should be used. Only a small minority of studies clearly provide measures of implicit and explicit interlimb^82,98,136^. Studies also vary in whether visual feedback is provided during the transfer phase. When feedback is present, transfer measures may be contaminated by additional learning with the untrained limb, whereas transfer blocks conducted without feedback provide a cleaner probe of how learning generalizes across limbs. Our meta-analysis only focused on the latter.

#### Primary outcome measures

The meta-analyses focused on three primary outcome measures. First, late adaptation indexes the adaptive changes in behavior measured during the last few trials of the perturbation block. This measure reflects the total amount of learning arising from both implicit and explicit processes. Second, we measured the aftereffect, assessed following removal of the perturbation (and in some cases removal of visual feedback). This measure provides a targeted measure of implicit adaptation. If a measure of late adaptation or aftereffect performance was defined in the study, we extracted this measure for our analysis. When these measures were not explicitly reported, late adaptation was defined as the final data point (trial or cycle) available in the adaptation time series, whereas aftereffects were defined as the first data point reported during the aftereffect phase.

Third, interlimb transfer was quantified as a percentage by dividing early performance during the transfer block with the untrained limb by late adaptation during the initial perturbation block achieved with the trained limb. When these phases were not explicitly defined in the original study, we operationalized late adaptation as the final datapoint (trial or cycle) in the reported time series and early transfer as the second datapoint in the transfer block, as the first trial can exhibit high variability due to context-switching effects when participants change limbs.

#### Effect size

For the learning-focused meta-analysis, the target effect size was Hedges’ g, a bias-corrected version of Cohen’s *d* that reduces small-sample bias and quantifies the standardized mean difference between the dominant and non-dominant hands^137^. For the interlimb transfer analysis, the target effect size was the transfer ratio (adaptation in the untrained limb divided by adaptation in the trained limb), which was natural log-transformed prior to analysis to improve distributional properties. For ease of interpretation, synthesized estimates and CIs were back-transformed and reported as transfer percentages.

Three independent coders (IN, AS, EM) extracted effect sizes from each study and discrepancies were resolved with consensus meetings. If the effect size was not reported, we calculated it using the F-statistic, t-statistic, or from the raw means and standard deviations extracted using WebPlotDigitizer (https://automeris.io/WebPlotDigitizer/). Because some samples contributed multiple effect sizes, violating the independence assumption of conventional random-effects meta-analysis, analyses were conducted using cluster-robust variance estimation^138^. Statistical significance was evaluated using small-sample correction, which applied Satterthwaite df to improve Type 1 error control when number of studies is limited^139^. Hedge’s g levels of 0.2, 0.5 and 0.8 correspond to small, moderate, and large effect sizes, respectively. Between-study heterogeneity was evaluated using I^2^ (ranges from 0 to 100%, with 25% representing low, 50% moderate and 75% high heterogeneity^140^. represents the between-study variance (its square root is equivalent to standard deviation). The meta-analysis was conducted in R using *metafor*^137^ for visualizations and *robumeta*^141^ for robust meta-regression tests. We used robust variance estimation to account for statistical dependence because 69.6% of the adaptation and 88% of the interlimb transfer studies reported more than one effect size.

#### Moderator analysis and publication bias

We used meta-regression to examine whether the effects of hand dominance and interlimb transfer varied as a function of task type (visuomotor vs. force-field adaptation), the number of targets, and the size of the perturbation (restricted to visuomotor rotation datasets). To assess publication bias, we conducted a robust Egger-type meta-regression test with *robumeta*, using the standard error of each effect size as a moderator.

### Experiments

#### Inclusion & Ethics Statement

All participants provided informed consent in accordance with policies approved by the Carnegie Mellon University’s Institutional Review Board. Participants completed Experiments 1-3 online in exchange for monetary compensation. All participants were recruited from Prolific, an online research recruitment platform.

#### Participants

We recruited a total of 526 healthy participants: 204 left-handed and 322 right-handed individuals. Hand preference was determined using a modified Edinburgh Handedness Inventory (Fig. S1) and used to screen for study eligibility. To qualify, participants had to report a *strong* preference for their dominant hand in activities of daily living, including writing, drawing, throwing, using the knife, and using the trackpad – the primary device used to complete the experiments online. For Experiments 1–2, participants first completed a speeded clicking task and a goal-directed reaching task to provide objective measures of dominant- and non-dominant-hand performance. They then proceeded to the main experimental task designed to assess motor learning.

#### Sample Size Justification

Experiments 1-3 were designed to explore the impact of hand dominance and handedness on implicit and explicit components of motor learning. As such, we determined the appropriate sample size using the data from the late adaptation meta-analysis. Studies reporting the strongest effects of hand dominance yielded effect sizes of approximately (= 0.25). A priori power analyses (α = 0.05, power = 0.80) indicated that a minimum of 40 participants per group would be required to detect small-to-moderate main effects. To ensure adequate power, we preregistered a target sample size of 50 participants per group. This yielded four groups in Experiments 1 and 2 (left-handed participants using their dominant or non-dominant hand and right-handed participants using their dominant or non-dominant hand), resulting in over 200 participants per experiment. Experiment 3 employed a comparable overall sample size but was restricted to right-handed participants.

We recognize that our final sample size exceeded the target specified in our preregistration (https://osf.io/jeqyf/overview). This deviation occurred because, once the study link was made available online, more participants than anticipated completed the experiment and were not excluded during our outlier removal procedure (Outlier Removal). We therefore retained these additional participants in our analysis, while ensuring that each group included at least the preregistered minimum of 50 participants (Table 1). This deviation does not affect our conclusions. Restricting the analyses to the first 50 participants per group who completed the study yielded the same pattern of results. Furthermore, repeating the analyses with the outlier participants included in Experiments 1–3 likewise revealed no evidence of asymmetries in either motor learning or interlimb transfer.

#### Apparatus

All participants used their own laptop to complete our study via a customized webpage. They used their computer trackpad to perform all tasks (sampling rate typically ∼60 Hz). Stimuli sizes automatically normalized to their screen sizes. The details below are provided as an example based on a 13” laptop screen with a resolution of 2560 x 1664 pixels.

#### Speeded-Clicking

Participants were instructed to click as quickly and accurately as possible on a blue target dot appearing at random locations on a 10-by-6 grid for 30 seconds per trial (Fig. 5A)^85^.

Participants completed six trials in total, alternating between their dominant and non-dominant hands, with the starting hand counterbalanced across participants. The first two trials served as familiarization and were not included in the analysis.

Upon a successful click, defined as clicking within the dot of a 1.45 x 1.45 mm, the target dot turned green for 0.5 seconds and then turned hollow again. After a 0.2-second delay, another blue dot would appear. Upon an unsuccessful click, the target turned red for 0.5 seconds, followed by the same delay before the next target appeared. We recorded both the total number of accurate clicks and response times per click, defined as time between the last and current click. Participants who were included in the main experiment were all included in the clicking assessment. For total clicks and response times, individual data points outside three standard deviations from the group mean were excluded from the analysis.

#### General Reaching Procedure

During each reaching trial, participants executed a planar movement from the center of the workspace to a visual target. The start position was indicated by a white circle (0.5 cm diameter) and the target by a blue circle (0.5 cm diameter). On a standard monitor, the radial distance between the start position and the target was 7 cm. Targets appeared at one of three locations (30°—upper right quadrant, 150°—upper left quadrant, 270°—lower center). Each cycle consisted of three reaches (one to each target location) presented in random order.

To initiate a trial, participants moved a white cursor (0.5 cm diameter) to the center start position. Visual feedback was only provided when the cursor was within 2 cm of the start position. Once the cursor was held at the start position for 500 ms, a target appeared, prompting participants to reach and ‘slice’ through the target. There were no constraints on reaction time. However, to discourage mid-movement corrections, the message ‘Too Slow’ was presented on the screen for 750 ms if the movement time exceeded 500 ms. To help guide the participant back to the start position, veridical feedback was provided when the hand moved back within 2 cm of the center.

#### Goal-directed Reaching

There were 20 cycles in total (60 trials). The experiment began with 10 movement cycles without visual feedback (30 trials): the first five cycles were performed with one hand and the next five with the other, with the starting hand (dominant vs. non-dominant) counterbalanced across participants. Veridical cursor feedback was then introduced for 10 additional cycles, again alternating between the dominant and non-dominant hands in the same order as the preceding block.

To assess movement precision, we calculated the standard deviation of endpoint movement angles when the movement crossed the target distance. We also recorded participants’ reaction time, defined as the interval between target onset and movement initiation, with movement initiation defined as the timepoint when the hand movement exceeded 1 cm. Both feedback and no-feedback blocks were averaged during analysis. Movement precision or reactions times that fell more than three standard deviations from the group mean and were excluded from the analysis for that specific variable.

#### Experiment 1: Implicit adaptation

To examine the lateralization of implicit motor learning, we used a clamped feedback task^74^, comparing the performance of four groups of participants: left- and right-handed individuals trained using either their dominant or non-dominant limb (Table 1). This task isolates implicit adaptation by providing non-contingent visual feedback that is spatially invariant with respect to the participant’s actual movement trajectory.

Participants completed a total of 132 cycles (396 trials). 10 baseline cycles without feedback (5 with the dominant hand, 5 with the non-dominant hand; order counterbalanced across participants), 10 baseline cycles with veridical feedback (5 with the dominant hand, 5 with the non-dominant hand; order counterbalanced across participants), 70 cycles with clamped feedback (using the randomly assigned trained limb), 10 cycles in a no-feedback aftereffect block (trained limb), 10 additional clamped-feedback cycles (trained limb), 10 cycles in an interlimb transfer block without feedback (switching to the untrained limb), 8 cycles in an interlimb learning block with clamped feedback (untrained limb), and 4 cycles in a no-feedback aftereffect block (untrained limb).

In the baseline no-feedback block, no cursor feedback was provided during the reaching movement. In the baseline veridical feedback block, the feedback cursor was aligned with the participant’s movement position, allowing the participants to become familiar with the basic reaching procedure and task requirements. In the perturbation block, we used clamped non-contingent feedback in which the radial position of the cursor was veridical, but the angular position followed an invariant trajectory, displaced by 30° relative to the target (the angular direction was counterbalanced across participants). To highlight the invariant nature of the clamped feedback, four demonstration trials were provided before the perturbation block. On all six trials, the target appeared straight down (270° position), and the participant was told to reach directly to the target; however, in each case the cursor was clamped at a different offset and direction from the target. In this way, the participants could see that the spatial trajectory of the cursor was unrelated to their own reach direction. In the no-feedback aftereffect block, the visuomotor rotation was removed, and feedback was not provided.

#### Experiment 2: Explicit Strategy

To examine the lateralization of explicit strategy use, we used a delayed feedback task^143^, comparing the performance of four groups of participants: left- and right-handed individuals trained using either their dominant or non-dominant limb (Table 1). This task isolates explicit adaptation by providing delayed endpoint visual feedback during learning—a manipulation shown to attenuate aftereffects and thereby bias performance toward explicit strategies.

Task schedule was very similar to that of Experiment 1: Participants completed a total of 98 cycles (298 trials). Ten baseline cycles without feedback (5 with the dominant hand, 5 with the non-dominant hand; order counterbalanced across participants), 10 baseline cycles with veridical feedback (5 with the dominant hand, 5 with the non-dominant hand; order counterbalanced across participants), 40 cycles with delayed feedback (using the randomly assigned trained limb), 10 cycles in a no-feedback strategy maintenance block (trained limb), 10 additional delayed feedback cycles (trained limb), 10 cycles in an interlimb transfer block without feedback (switching to the untrained limb), and 8 cycles in a no-feedback maintain block (untrained limb).

Veridical and no-feedback phases were identical to those in Experiment 1. The key manipulation in this experiment occurred during the delayed feedback phase, in which cursor feedback rotated by rotated by 60° in either the clockwise or counterclockwise direction (counterbalanced across participants). Endpoint cursor feedback was delayed by 800 ms, a manipulation shown to attenuate implicit adaptation and thereby bias performance towards strategy use.

#### Experiment 3: Continuous Tracking

To examine hemispheric specialization in a more ecological form of motor learning, we employed a continuous manual tracking task, comparing two groups of right-handed participants who performed the task with either their dominant or non-dominant hand (Table 1). Participants used a computer mouse to continuously track a moving target displayed on the screen. To generate a smooth yet unpredictable trajectory and minimize opportunities for sequence learning, the target’s motion was constructed by summing 14 independent sine waves (seven along the X-axis and seven along the Y-axis). The component frequencies ranged from 0.05 to 1.075 Hz, producing a fundamental period of 40 s. Because each trial lasted only 30 s, the target trajectory never repeated within a trial, preventing participants from anticipating its future path.

The experimental design mirrored that of the previous experiments and consisted of 56 trials divided into four phases. During Baseline (16 trials), participants tracked the target with veridical feedback. During Early Adaptation (30 trials), a visuomotor rotation was gradually introduced at a rate of 1° per trial until reaching 30°. The perturbation was then held constant during Late Adaptation (6 trials) to assess asymptotic performance. During Washout (4 trials), the rotation was abruptly removed and veridical feedback reinstated to measure aftereffects.

### Data Analysis

All data processing and statistical analyses were conducted in R. The primary dependent measure was movement angle. In Experiments 1–2, movement angle was defined as the angular position of the hand at the moment it reached the radial target distance of 7 cm from the start position. In Experiment 3, movement angle was computed continuously for each sampled frame throughout the tracking trajectory.

Movement angles were baseline-corrected by subtracting each participant’s mean movement angle during the baseline phase from all subsequent movement angles. For counterclockwise rotations, movement angles were sign-flipped (multiplied by −1) so that positive deviations consistently reflected learning in the direction opposite the perturbation (i.e., the intended compensatory direction).

Several learning phases were defined a priori. Early adaptation was quantified as the mean movement angle across the first ten movement cycles following introduction of the rotation (cycles 20–30 in Experiments 1 and 2; trials 15–25 in Experiment 3). Late adaptation was quantified as the mean movement angle across the final ten movement cycles of the rotation block (cycles 80–90 in Experiment 1, cycles 50–60 in Experiment 2, and trials 45–55 in Experiment 3). Aftereffects and washout were quantified as the mean movement angle across the entire no-feedback or washout block (aftereffects: cycles 90–100 in Experiment 1; washout: trials 50–55 in Experiment 3). Interlimb transfer was quantified as the mean movement angle across the transfer block (cycles 110–120 in Experiment 1; cycles 80–90 in Experiment 2).

These data were analyzed using a linear mixed-effects model (R function: lmer)^144^, with handedness group (left vs. right) and hand dominance (dominant vs. non-dominant) included as fixed effects, and participant ID included as a random effect. Main effects were evaluated using Type III analysis of variance tests (R function: Anova)^145^. Post hoc comparisons between groups and conditions were conducted (R function: emmeans)^146^.

#### Outlier Removal

Exclusion criteria varied between experiments due to considerations regarding the learning process elicited. For Experiment 1, we excluded reaching trials in which the movement angle deviated from a 10-trial trendline by more than 3 standard deviations. These datapoints are likely to reflect attentional lapses, since participants were instructed to reach directly to the target. This resulted in the exclusion of 1.5% [1%, 3.2%] (Median [IQR]) of the trials.

Outlier participants were defined using two criteria: (1) participants for whom outlier trials defined above exceeded 3.7% of the experiment, and (2) participants whose baseline movement precision (standard deviation of movement angle) exceeded 40° (a value far exceeding typical baseline motor variability: ∼5–10°). We collected data from 274 online participants to obtain a final sample of 202 participants (N = ∼50 per group, Table 1).

For Experiment 2, we sought to determine whether strategy use is lateralized to a specific hemisphere. Accordingly, we restricted our sample to individuals who clearly deployed a re-aiming strategy— operationalized as movement angles exceeding 10° during late adaptation and no-feedback maintenance trials (assessed via one-sample t-tests). To obtain a final sample of at least 217 participants (N = ∼50 per group, Table 1), we collected data from 627 online participants. We recognize that the resulting exclusion rate is relatively high; however, this is a strength of the design, as it enables a more conservative test of our primary hypothesis by excluding participants who did not reliably engage a strategy.

For Experiment 3, outliers were identified and excluded at both the trial and participant levels. Trial-level exclusions were applied when angular error exceeded 1.5 × the interquartile range (IQR) of the group distribution for a given trial, resulting in the removal of 3.8% of all observations. Participant-level exclusions were applied according to two predefined criteria: (1) incomplete datasets or (2) failure to provide valid trial data due to technical issues, such as missing database entries or failed session initialization. Overall, 33 of the 140 enrolled participants were excluded according to the predefined criteria, resulting in a final sample of 107 participants for analysis.

#### Cluster-based permutation test

We used a cluster-based permutation test to identify clusters of cycles in which movement angle (Fig. 7, 8, and 9) and movement kinematics (reaction time, movement time, and search time; Fig. S2, S3, and S5) differed across groups^75,147,148^. The test comprised of two steps. First, a F test (comparing all four experimental conditions) was performed on each movement cycle in the perturbation block across experimental conditions to identify cluster that showed a significant difference. Clusters were defined as epochs in which the P value from the F tests was less than 0.05 for at least two consecutive trials. The F values were then summed up across cycles within each cluster, yielding a combined cluster score. Second, to assess the probability of obtaining a cluster of consecutive cycles with significant p-values, we performed a permutation test. Specifically, we generated 100,000 permutations by shuffling the condition labels. For each shuffled permutation, we computed the sum of F scores to generate a null distribution of scores. The proportion of random permutations that resulted in a F score that was greater than or equal to that obtained from the data could be directly interpreted as the P value. Clusters with P_perm_ < 0.05 are reported. Post hoc t-tests were conducted on significant clusters using 100,000 permutations to assess pairwise group differences.

#### Model-based Analysis

To estimate the learning rate during motor learning, we fit a standard state-space model to each participant’s reaching data. The model assumes that behavior is driven by an internal state *x_n_*, which reflects the participant’s current estimate of the perturbation. Overall performance on trial *n* (the predicted reach angle *y_n_*) reflects this internal state plus motor noise:

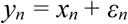

where *ε_n_* ∼ *N(0, σ_M_)* is motor variability with standard deviation *σ_M_*.

The internal state is updated incrementally from trial to trial as a function of two parameters: the retention factor *A*, which determines how much of the state is retained across trials, and the learning rate *B*, which governs how strongly errors drive updating.

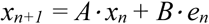

The definition of the error signal *e_n_* differs across experiments. In Experiments 1, participants were instructed to reach directly to the target and ignore the clamped feedback which was rotated relative to the target location. Under these conditions, the visual feedback remained invariant across trials and was defined as the imposed clamped visuomotor rotation *r*:

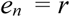

Thus, retention *A* controls memory of the previous trial, and learning rate *B* determines sensitivity to error.

In Experiments 2, the participants’ objective was to counteract a visuomotor perturbation. In the extreme, this would mean aligning the visually perturbed cursor, rotated with respect to the participant’s movement angle, with the target. Accordingly, error was defined as the difference between the imposed perturbation *r* and the current internal state *x_n_*:

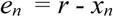

We fit the model to each participant’s reaching data during the adaptation block by minimizing the negative log-likelihood of the observed reach angles given the model predictions. The optimization was carried out in R using the optim() function with the “L-BFGS-B” method and 10 random starting points. The three free parameters were the retention factor A, the learning rate B, and motor noise *σ_M_*. Bounds were set as follows: A ∈ [0.001,1], B ∈ [0.001,1], and σ ∈ [0.001,100]. The best-fitting parameter B was taken as the participant’s learning rate estimate.

## Supplementary Figures

**Figure S1.**
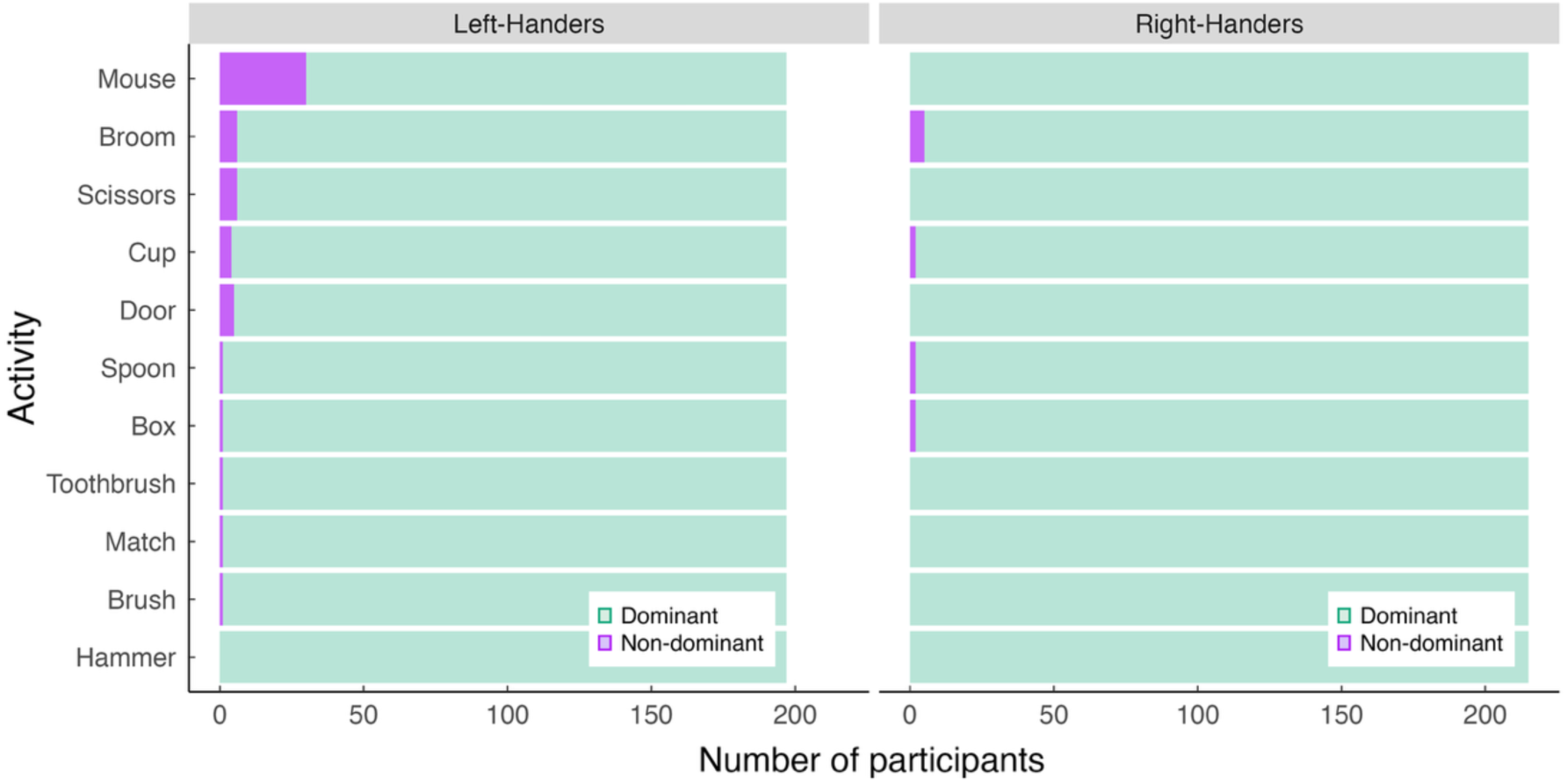
Participant pool exhibited strong left- and right-hand dominance. Number of participants reporting a preference for using their dominant (green) or non-dominant (purple) hand for each item on the modified Edinburgh Handedness Inventory. Participants were included in the study only if they identified their dominant hand as their preferred hand for writing, drawing, throwing, using a trackpad, and using a knife.

**Figure S2.**
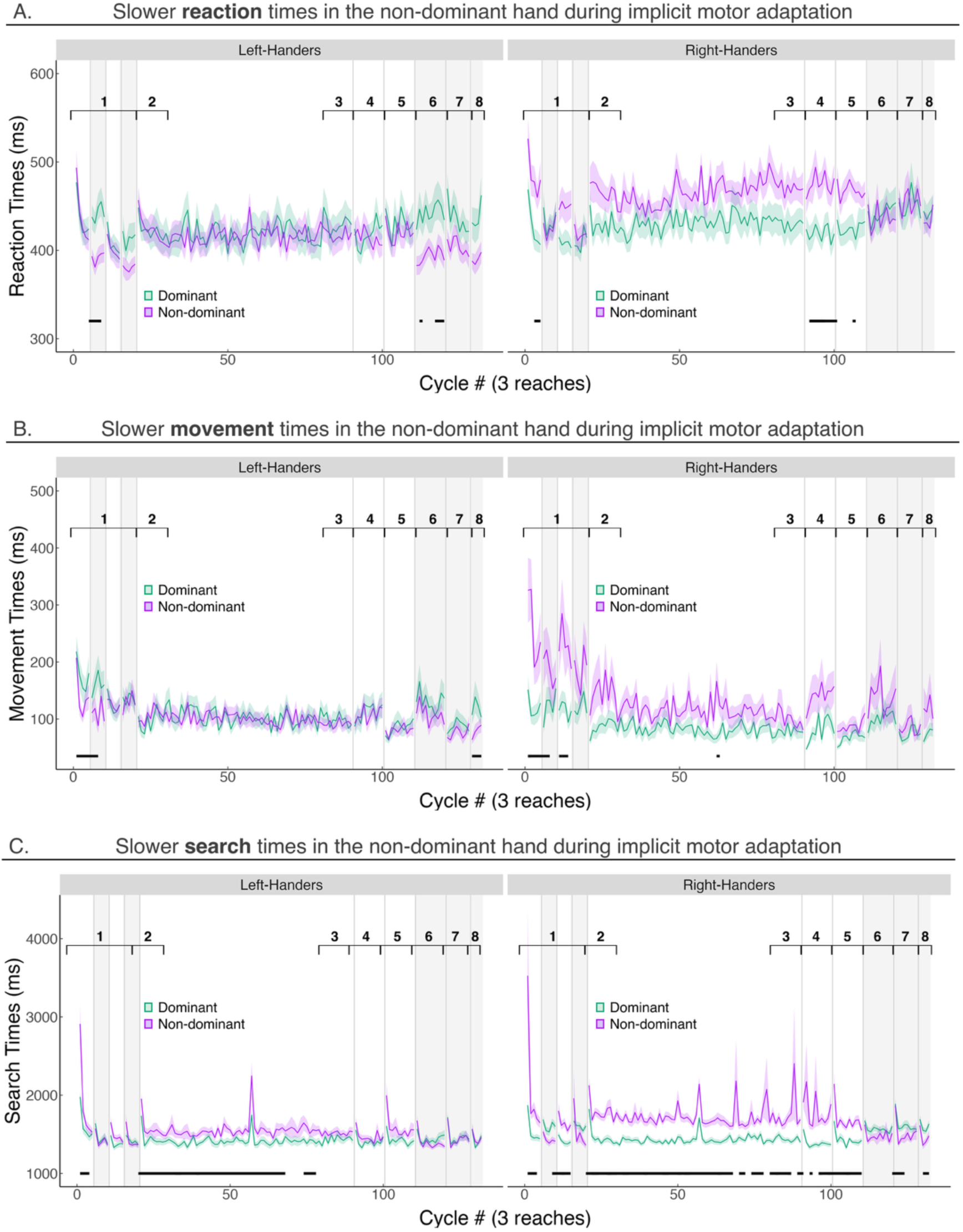
Movement kinematics in Experiment 1. **A–C)** Mean reaction time (interval between target appearance and movement initiation, defined as moving 1 cm), movement time (interval between movement initiation and reaching the 7-cm target distance, marking movement termination), and search time (interval between movement termination and returning to the start position) for participants trained with the dominant or non-dominant hand across task phases (black numbers). The green curve denotes the group trained with the dominant hand, whereas the purple curve denotes the group trained with the non-dominant hand. Light gray bars indicate blocks in which participants temporarily switched to their untrained hand. Black horizontal lines at the bottom of each panel denote clusters of significant differences (p < 0.05) between the dominant- and non-dominant-hand groups (between-participant analysis), identified using cluster-based permutation tests. Note that in Fig. 6F, reaction times for the dominant and non-dominant hands were compared using a within-participant analysis of the baseline condition (block 1), revealing a robust dominant-hand advantage.

**Figure S3.**
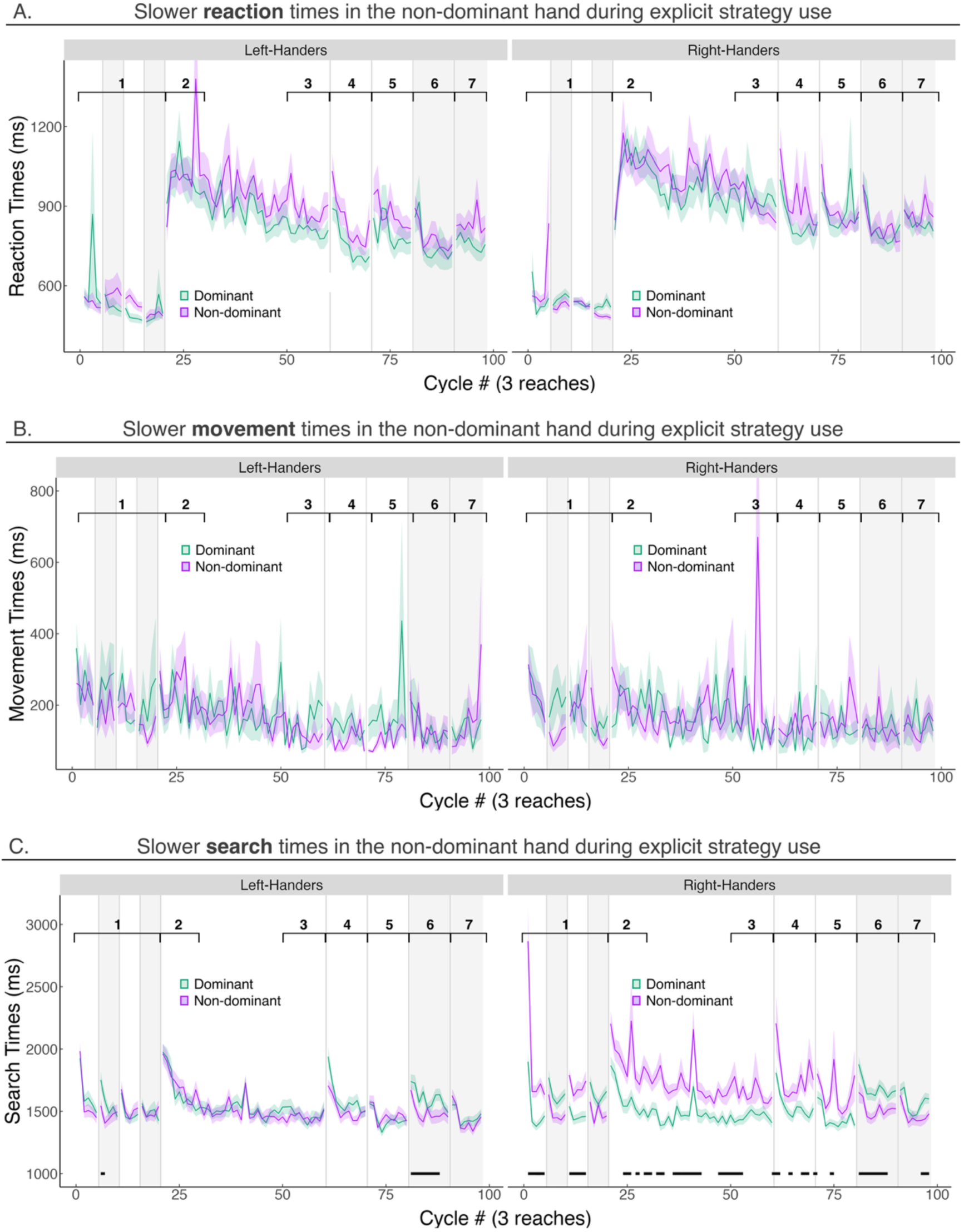
Movement kinematics in Experiment 2. **A–C)** Mean reaction time (interval between target appearance and movement initiation, defined as moving 1 cm), movement time (interval between movement initiation and reaching the 7-cm target distance, marking movement termination), and search time (interval between movement termination and returning to the start position) for participants trained with the dominant or non-dominant hand across task phases (black numbers). The green curve denotes the group trained with the dominant hand, whereas the purple curve denotes the group trained with the non-dominant hand. Light gray bars indicate blocks in which participants temporarily switched to their untrained hand. Black horizontal lines at the bottom of each panel denote clusters of significant differences (p < 0.05) between the dominant- and non-dominant-hand groups, identified using cluster-based permutation tests. Note that in Fig. 6F, reaction times for the dominant and non-dominant hands were compared using a within-participant analysis of the baseline condition (block 1), revealing a robust dominant-hand advantage.

**Figure S4.**
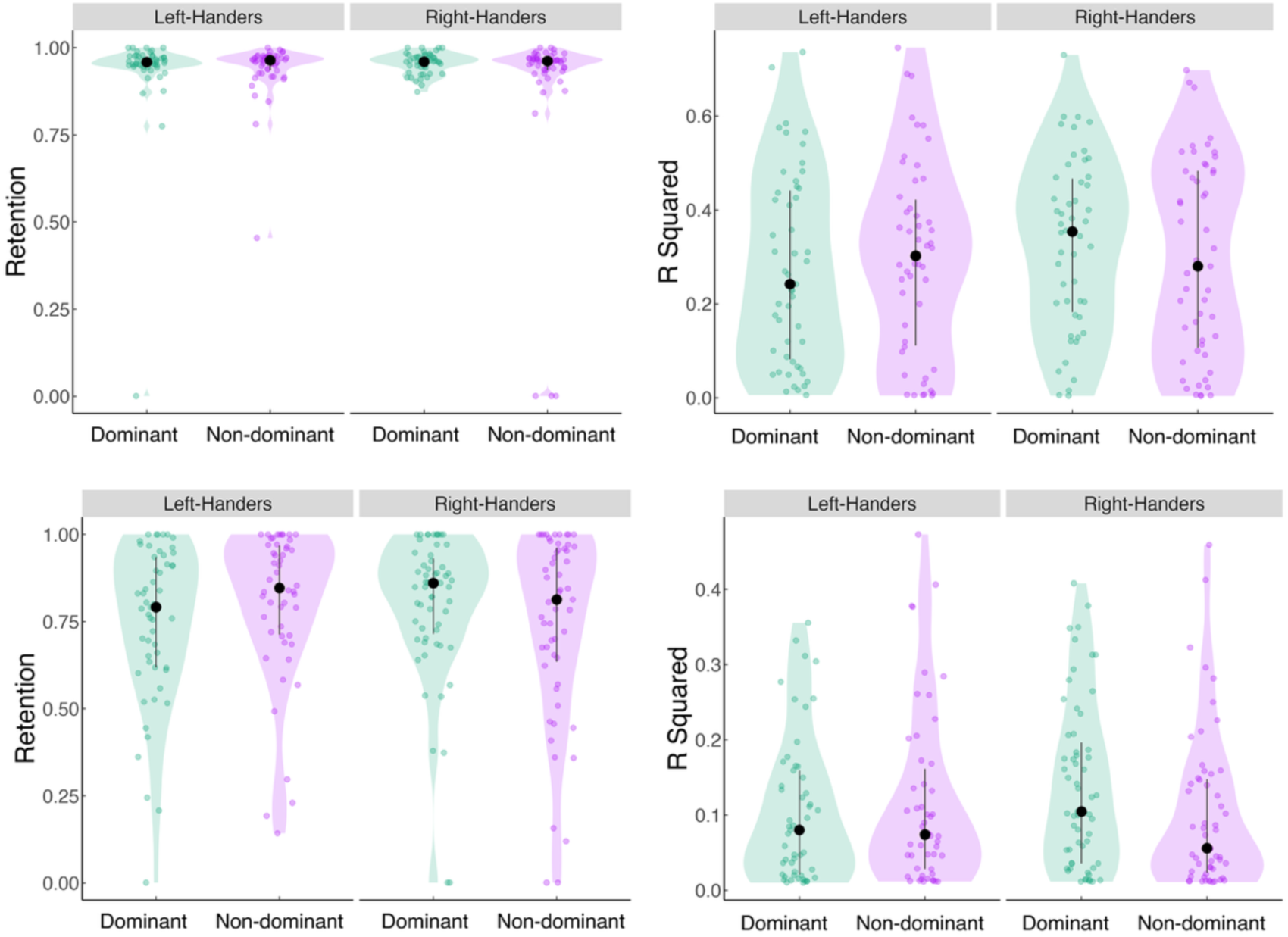
State-space model parameters and goodness of fit. **A)** Retention parameter and **B)** goodness of fit (R²) obtained by fitting the state-space model to the data from Experiment 1 (top row) and Experiment 2 (bottom row). Learning rates are shown in the main text. Solid circles denote the group median, error bars indicate the interquartile range, and translucent dots represent individual participants. Violin widths reflect the underlying data density. No significant differences were observed between the dominant- and non-dominant-hand groups for either measure.

**Figure S5.**
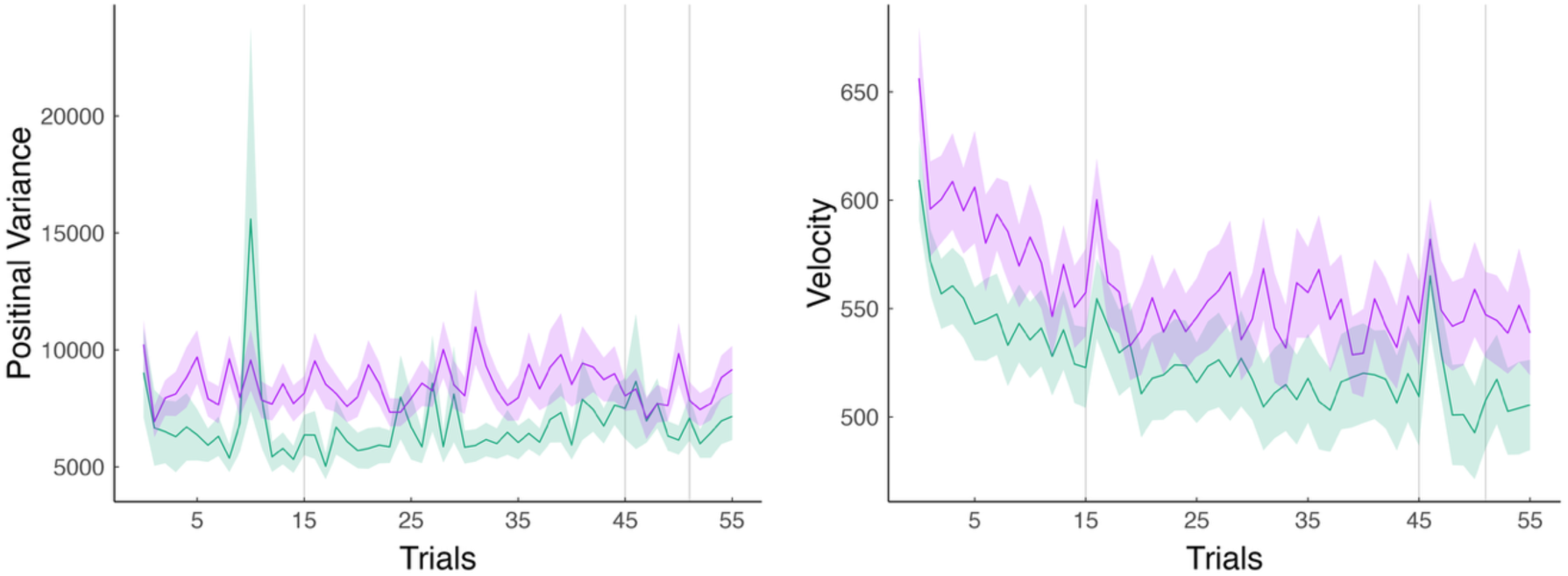
Movement kinematics in Experiment 3. Position variance and mean movement velocity for participants trained with the dominant or non-dominant hand. green curve denotes the group trained with the dominant hand, whereas the purple curve denotes the group trained with the non-dominant hand.

## Supplementary Methods

### Search Terms for Meta-analysis

Our search terms for the adaptation meta-analysis were (handedness OR hand dominance OR lateralization OR hemispheric differences OR bilateral OR effector-dependent OR limb-dependent OR effector-independent OR limb-independent AND motor adaptation OR sensorimotor adaptation OR visuomotor adaptation OR motor learning OR perturbation). We also included studies identified to us by other researchers.

Our search terms for the interlimb transfer meta-analysis were (text: interlimb OR intermanual OR lateralization OR hemispheric differences OR bilateral OR effector-dependent OR limb-dependent OR effector-independent OR limb-independent AND text: transfer OR motor adaptation OR sensorimotor adaptation OR visuomotor adaptation OR motor learning OR perturbation). We also included studies identified to us by other researchers.

